# Near-atomic structures of the BBSome reveal a novel mechanism for transition zone crossing

**DOI:** 10.1101/2020.01.29.925610

**Authors:** Kriti Bahl, Shuang Yang, Hui-Ting Chou, Jonathan Woodsmith, Ulrich Stelzl, Thomas Walz, Maxence V. Nachury

**Author notes:** Department of Therapeutic Discovery, Amgen Inc., South San Francisco, CA 94080, USA. These authors contributed equally to this work and are listed alphabetically (Kriti Bahl, Shuang Yang).

## Abstract

The BBSome is a complex of eight Bardet-Biedl Syndrome (BBS) proteins that removes signaling receptors from cilia. The GTPase ARL6/BBS3 recruits the BBSome to the ciliary membrane where the BBSome–ARL6^GTP^ complex ferries G protein-coupled receptors (GPCRs) across the transition zone, a diffusion barrier at the base of cilia. Here, we find that the BBSome undergoes a conformational change upon recruitment to membranes by ARL6^GTP^. Modeling the binding of the BBSome to membranes and to the GPCR Smoothened (SMO) reveals that the amphipathic helix 8 of SMO must be released from the membrane for SMO to be recognized by the BBSome. Underscoring the functional importance of amphipathic helix extraction in TZ crossing, we find that exchanging the amphipathic helix of ARL6 for one that embeds deeper into the membrane blocks BBSome-mediated exit of GPCRs from cilia. We propose that the rigid curvature and dense lipid packing of the transition zone reject asymmetric insertions in the inner leaflet and that the BBSome licenses transition zone crossing by extracting bulky amphipathic helices from the inner leaflet.

## INTRODUCTION

Cilia dynamically concentrate signaling receptors to sense and transduce signals as varied as light, odorant molecules, Hedgehog morphogens and ligands of G protein-coupled receptors (GPCRs) (Anvarian et al., 2019; Bangs and Anderson, 2017; Nachury and Mick, 2019). Highlighting the functional importance of dynamic ciliary trafficking, the appropriate transduction of Hedgehog signal relies on the disappearance of the GPCR GPR161 and the Hedgehog receptor Patched 1 from cilia and the accumulation of the GPCR Smoothened (SMO) within cilia (Bangs and Anderson, 2017). Regulated exit from cilia represents a general mechanism to redistribute signaling molecules on demand (Nachury and Mick, 2019). Patched 1, GPR161, SMO and other ciliary membrane proteins are all ferried out of cilia in a regulated manner by an evolutionarily conserved complex of eight Bardet-Biedl Syndrome (BBS) proteins, the BBSome (Nachury, 2018; Wingfield et al., 2018). While GPR161 and other ciliary GPCRs such as the Somatostatin receptor 3 (SSTR3) are removed from cilia by the BBSome only when they become activated, SMO undergoes constitutive BBSome-dependent exit from cilia in unstimulated cells to keep its ciliary levels low. Accumulation of SMO in cilia is then, at least in part, achieved by suppression of its exit (Milenkovic et al., 2015; Nachury and Mick, 2019; Ye et al., 2018).

Membrane proteins travel into, out of, and within cilia without utilizing vesicular intermediates and remain within the plane of the ciliary membrane (Breslow et al., 2013; Chadha et al., 2019; Milenkovic et al., 2009; Ye et al., 2018). Thus, membrane proteins that enter and exit cilia must cross the transition zone (TZ), a diffusion barrier at the base of cilia, by lateral transport (Garcia-Gonzalo and Reiter, 2017). How the TZ lets privileged membrane proteins across while blocking the diffusion of most membrane proteins remains an open question. Recently, we found that regulated TZ crossing of GPR161 is enabled by the BBSome in concert with the ARF-like GTPase ARL6/BBS3 (Ye et al., 2018), and two models have been proposed for BBSome/ARL6- mediated passage through the TZ (Nachury and Mick, 2019). The first model takes into account the molecular motors that move intraflagellar transport (IFT) trains up (anterograde movement) and down (retrograde movement) the ciliary shaft and the association of the BBSome with IFT trains to propose that the retrograde motor dynein 2 physically drags the BBSome–ARL6^GTP^– IFT-B complex with its associated GPCRs across the TZ (Nachury, 2018; Wingfield et al., 2018). Alternatively, the membrane-bound BBSome may grant its attached cargoes a license to cross the TZ, in analogy to how karyopherins have been proposed to allow cargoes to permeate the nuclear pore complex interior (Schmidt and Görlich, 2016). In the absence of a direct test for these hypotheses and considering that other models may exist, the mechanism of facilitated TZ crossing by the BBSome remains a fundamental unanswered question.

Our recent cryo-electron microscopy (cryo-EM) structure of the BBSome revealed that the BBSome exists mostly in an auto-inhibited, closed conformation in solution and undergoes a conformational change as it is recruited to membranes by ARL6^GTP^ (Chou et al., 2019). Given that ARL6^GTP^ triggers polymerization of a membrane-apposed BBSome/ARL6 coat (Jin et al., 2010) and enables BBSome-mediated TZ crossing (Ye et al., 2018), the ARL6^GTP^-bound BBSome conformation represents the active form of the complex. Here we determine high-resolution structures of the BBSome alone and bound to ARL6^GTP^, and we map the BBSome–SMO interaction to model how the membrane-associated BBSome–ARL6^GTP^ complex recognizes its cargoes. Surprisingly, our studies reveal that SMO must eject its amphipathic helix 8 (SMO^H8^) from the inner leaflet of the membrane in order to be recognized by the BBSome. Based on the known curvature and lipid composition of the TZ, we propose that extraction of SMO^H8^ through its interaction with the BBSome licenses SMO for TZ crossing and we present functional studies in support of this model.

## RESULTS

### High-resolution structure and model of the BBSome

Following on our previous strategy (Chou et al., 2019), we purified the BBSome to near-homogeneity from retinal extract and analyzed its structure by single particle cryo-electron microscopy. The advent of higher throughput direct detector cameras and faster automated data-collection procedures combined with improvements in data processing with new tools implemented in RELION-3 (Zivanov et al., 2018) led to a BBSome map at an overall resolution of 3.4 Å from an initial dataset of 770,345 particles (**Fig. S1, S3A**).

The BBSome is composed of 29 distinct domains characteristic of sorting complexes (**Fig. 1A**). α-solenoids, β-propellers, pleckstrin homology (PH) and appendage domains are all present in multiple copies and our previous map made it possible to build a Cα backbone model that encompassed 25 out of 29 domains (PDB-Dev accession PDBDEV_00000018; Chou et al., 2019). For building the current model, the previous Cα model was docked into the map, and the higher-resolution map enabled us to confidently assign side chains for most regions (**Fig. 1B**).

**Figure 1.**
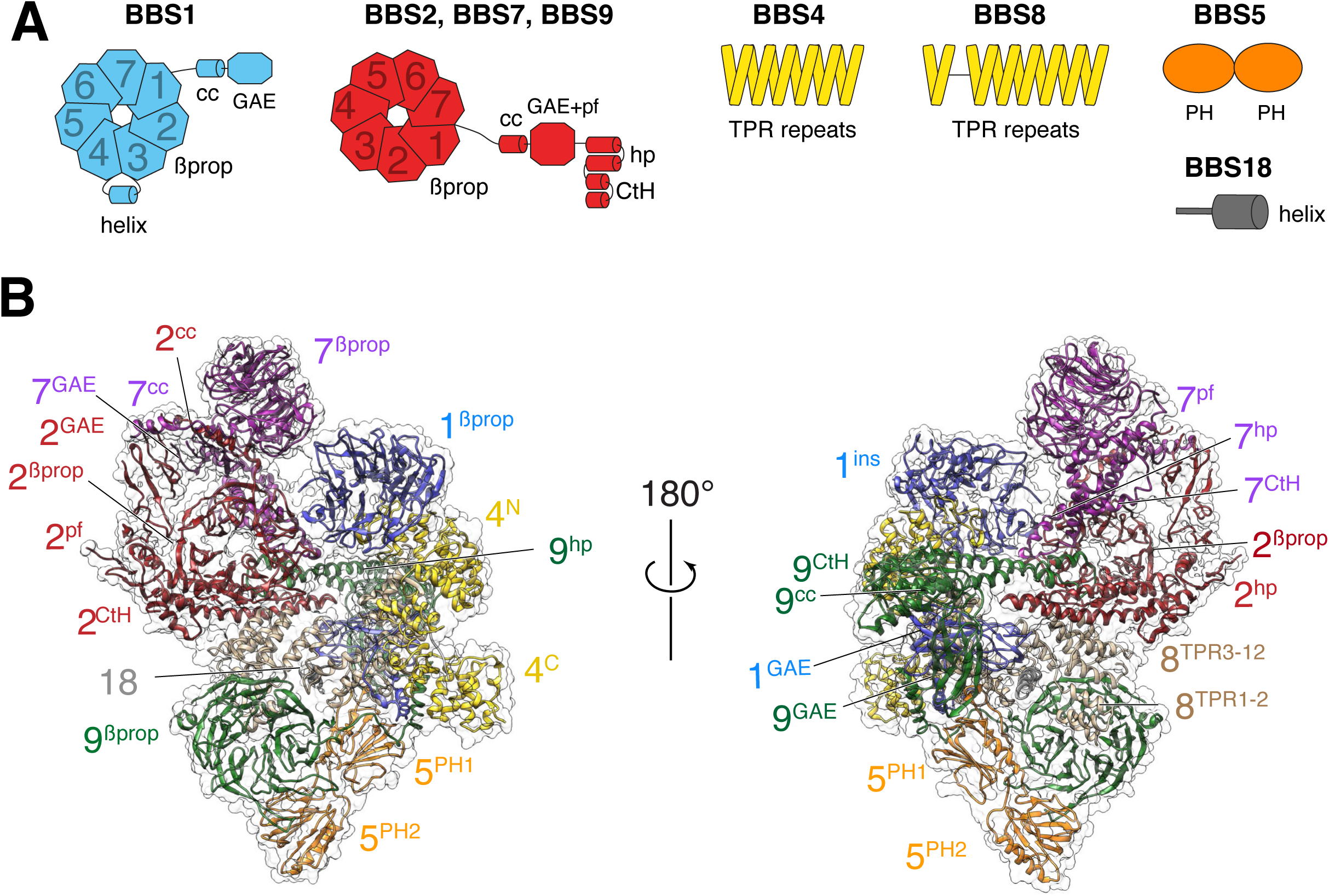
Overall structure of the BBSome. **A.** Diagrams showing the domain architecture of the eight BBSome subunits. βprop, β-propeller; cc, coiled coil; GAE, γ-adaptin ear; pf, platform; ins, insert; hp, hairpin; CtH, C-terminal helix bundle; TPR, tetratricopeptide repeat; PH, pleckstrin homology. **B.** Two views of the cryo-EM map (transparent surface) and the near-atomic model of the BBSome complex shown in ribbon representation. Individual domains are labeled with the numbers identifying the subunit and the superscripts denoting the specific domain.

The new map allowed us to build the coiled-coil domains of BBS1 and BBS9, for which densities were not well-defined in the previous map. Altogether, 27 out of 29 domains distributed across the 8 BBSome subunits could be modeled. Despite the increased resolution of the current density map, however, the gamma-adaptin ear (GAE) domains of BBS2 and BBS7 could not be modeled, and side chains could not be assigned for BBS2^βprop^, BBS2^cc^, BBS7^βprop^ and BBS7^cc^.

### High-resolution structure of the BBSome bound to ARL6^GTP^

Consistent with our previous observations based on a 4.9-Å resolution map of the BBSome (Chou et al., 2019), the new BBSome structure cannot accommodate binding to ARL6^GTP^. Fitting a homology model of the bovine BBS1^βprop^–ARL6^GTP^ complex (based on the X-ray structure of the *Chlamydomonas* complex; Mourão et al., 2014) in either BBSome structure caused a steric clash between ARL6^GTP^ and a region encompassing BBS2^βprop^ and BBS7^cc^. These data support a model in which the BBSome exists in an autoinhibited form in solution and undergoes a conformational opening upon recruitment to membranes by ARL6^GTP^, similar to other sorting complexes such as COPI, AP-1 and AP-2 (Cherfils, 2014; Faini et al., 2013).

The membrane-associated form of the ARL6^GTP^-bound BBSome represents its active conformation, because ARL6^GTP^ enables TZ crossing (Ye et al., 2018). To determine the nature and consequence of the conformational change in the BBSome that takes place upon ARL6^GTP^ binding, we set out to determine the structure of the BBSome–ARL6^GTP^ complex.

Mixing recombinant ARL6^GTP^ together with the purified BBSome allowed for complex formation in solution. The BBSome–ARL6^GTP^ complex was analyzed by cryo-EM (**Fig. S2A, B**), yielding a density map at an overall resolution of 4.0 Å (**Fig. S3A**). Focused refinement of the top and lower lobes of the complex resulted in improved maps of 3.8 Å and 4.2 Å resolution, which facilitated model building (**Fig. 2A and S2C**). Even though the apparent overall resolution was nominally not as good as that of the BBSome alone, several domains were better resolved in the density map of the BBSome–ARL6^GTP^ complex (**Fig. S3C**). In particular, the quality of the map was significantly increased for the top β-propeller (**Fig. S3B**). The improved map quality allowed us to correctly place the β-propellers (βprop) of BBS2 and BBS7, which were swapped in our previous structural description (Chou et al., 2019) due to their extreme similarity and the limited resolution of the previous map. This new assignment is further supported by a recently published structure of the BBSome (Singh et al., 2020).

**Figure 2.**
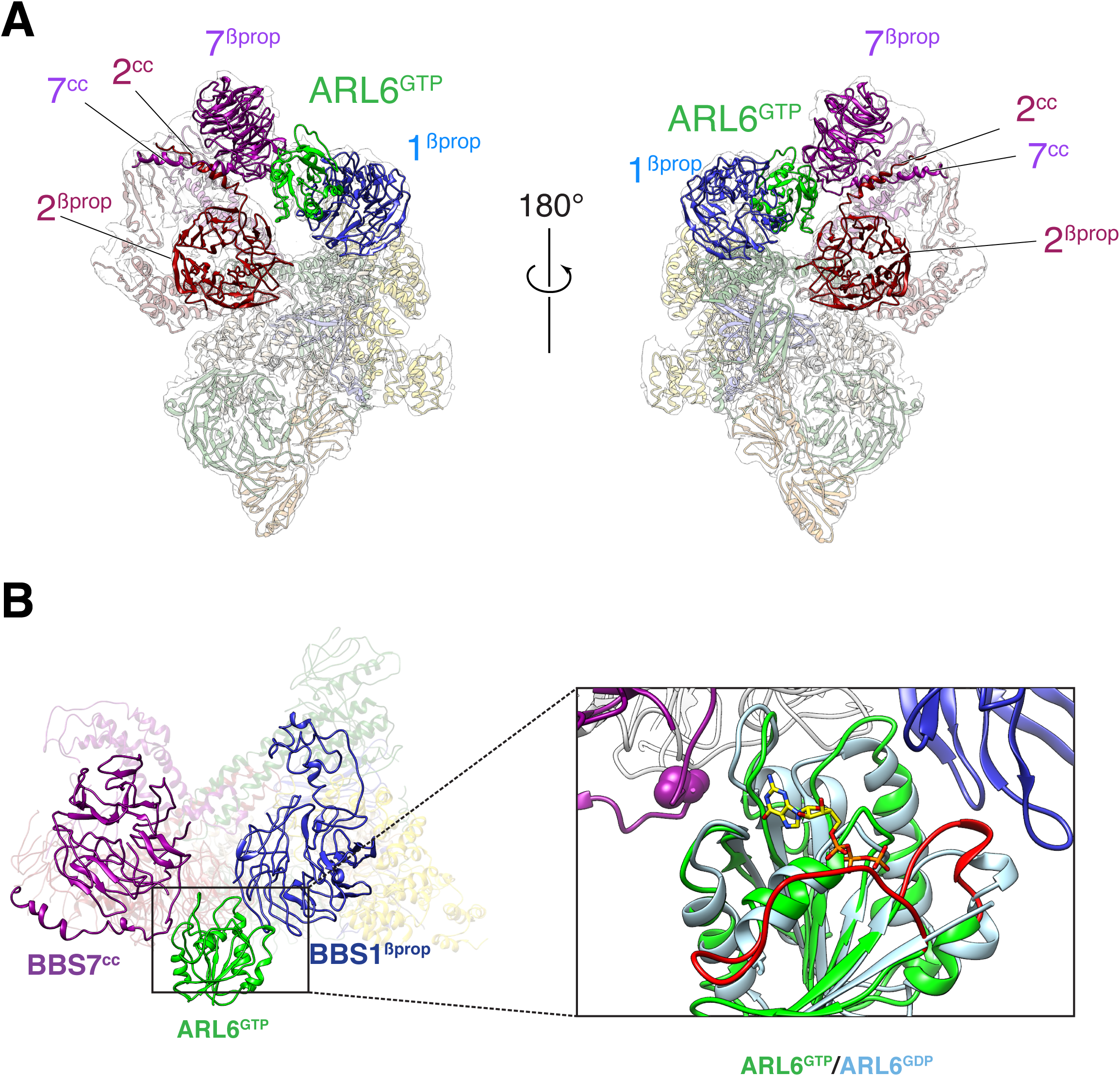
Overall structure of the BBSome–ARL6^GTP^ complex. **A.** Two views of the cryo-EM map (transparent surface) and the near-atomic model of the BBSome–ARL6^GTP^ complex shown in ribbon representation. BBSome domains that move as a result of ARL6^GTP^ binding are highlighted and labeled. **B.** Overall view (left panel) of the BBSome–ARL6^GTP^ complex and close-up view (right panel) focusing on the interaction of ARL6^GTP^ with the loop connecting BBS7^cc^ and BBS7^βprop^, and with BBS1^βprop^. As previously shown (Mourão et al., 2014) and consistent with other small GTPases, ARL6^GTP^ contacts the BBS1 β-propeller with its Switch1 and Switch2 regions (red) that change conformation upon exchange of GDP for GTP. In contrast, ARL6 contacts BBS7 using a surface that is largely unaffected by nucleotide binding (light blue for ARL6^GDP^ and lime green for ARL6^GTP^). A homology model of bovine GDP-bound ARL6^GTP^ (based on the crystal structure of the *Chlamydomonas* protein; PDB ID: 4V0K) was aligned to the model of the GTP-bound ARL6 in our BBSome–ARL6^GTP^ complex. The Gly329 residue mutated to Val in a BBS patient is depicted as magenta atom balls.

In the BBSome–ARL6 structure, ARL6^GTP^ is nestled in a wedge opening between BBS1^βprop^ and BBS7^βprop^. A ∼20° rotation of BBS1^βprop^ from the BBSome alone conformation allows ARL6^GTP^ to move away from the steric clash with BBS2^βprop^. This movement of BBS1^βprop^ is accompanied by a twisting of the first two TPR repeats from the BBS4 α-solenoid (**Video 1**), in line with the close association between the N terminus of BBS4 and BBS1^βprop^ seen in the BBSome alone structure and confirmed by cross-link mass spectrometry (Chou et al., 2019). Besides the movements of BBS1^βprop^ and BBS4^TPR1-2^, ARL6^GTP^ binding caused only subtle changes in the structure of the BBSome. The movements of BBS4 and BBS1 are in agreement with two recently published structures of the ARL6^GTP^-bound BBSome (Klink et al., 2020; Singh et al., 2020).

We note that the conformational opening of the BBSome is likely spontaneous as a minor 3D class corresponding to the open form could be detected in our previous data set of the BBSome alone. As the 3D class of the open conformation contained only a small percentage of the particles in the data set, the equilibrium between closed and open form in solution is strongly shifted towards the closed form. Binding to ARL6^GTP^ would thus act as a thermodynamic sink that locks the BBSome into the open conformation.

Small GTPases of the ARF/ARL family undergo conformational changes in 3 regions upon nucleotide exchange from GDP to GTP: the Switch 1 and 2 regions and the Interswitch toggle (Sztul et al., 2019). As previously found in the crystal structure of the BBS1^βprop^–ARL6^GTP^ complex (Mourão et al., 2014), the BBS1^βprop^ makes contacts with the Switch 2 region and with helix *α*3 of ARL6^GTP^ while the Switch 1 region of ARL6 is readily available for interacting with other, yet unidentified, complexes (**Fig. 2B**). Interestingly, the ‘backside’ of ARL6^GTP^ (i.e., the surface on the opposite side of the Switch regions) interacts with a loop that connects BBS7^βprop^ and BBS7^cc^. Given the absence of conformational changes in the backside of ARL6 upon nucleotide exchange, ARL6 binding to the BBS7^βprop^-BBS7^cc^ loop will not be gated by the nucleotide state, similar to the proposed binding of ARF1 to the γ subunit of the clathrin adaptor AP-1 (Ren et al., 2013). The homozygous mutation of the Gly329 residue to Val in a BBS patient (Chou et al., 2019) is predicted to interfere with the binding of ARL6 backside to BBS7 (**Fig. 2B**), suggesting functional importance for this interaction.

In addition, the physical interaction between ARL6^GTP^ and the upper lobe removes the upper lobe flexibility previously observed in the BBSome alone preparation, resulting in a more stable BBSome conformation (**Fig. 3B**).

**Figure 3.**
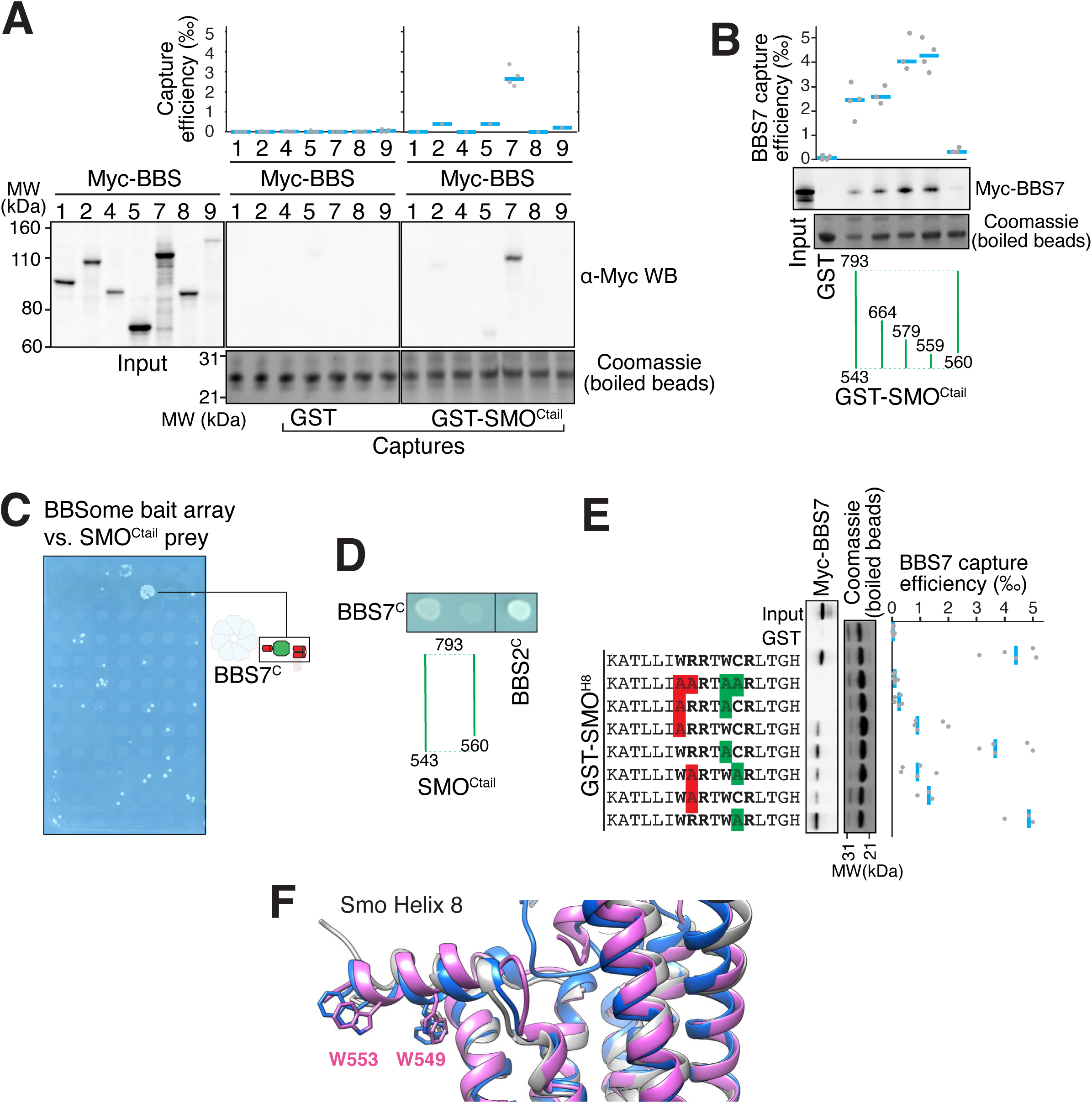
The BBSome recognizes SMO via membrane-embedded residues in SMO helix 8. **A-B.** GST-capture assays were conducted with *in vitro* translated BBSome subunits tagged with a 6xMyc epitope. Bound material was released by specific cleavage between GST and the fused peptide, and released proteins were detected using a Western blot and anti-Myc antibody (*α*-Myc WB). The proportions of BBSome subunits recovered in the eluate are plotted; grey circles are individual data points and blue lines are mean values. Even loading of the glutathione beads is demonstrated by staining for the remaining GST-tagged proteins after cleavage elution. **A.** Capture of individual BBSome subunits with GST-SMO^Ctail^ (aa 543-793) identifies BBS7 as the SMO-binding subunit. **B.** Capture assays with truncations of SMO^Ctail^ find that SMO^H8^ is necessary and sufficient for binding to BBS7. **C.** Yeast two-hybrid (YTH) assays with SMO^Ctail^ against an array of BBS protein fragments identify an interaction between a C-terminal fragment of BBS7 (BBS7^C^, residues 326-672) and SMO^Ctail^. The composition of the BBS YTH array is shown in Table S2. **D.** YTH assays find that SMO^H8^ is required for the interaction with BBS7^C^. Growth controls on diploid-selective medium for panels **C-D** are shown in **Fig. S4C**. **E.** Capture assays of BBS7 with mutants of SMO^H8^ (aa 543-559) identify Trp549 and Trp553 as the major BBS7-binding determinants of SMO^H8^. **F.** Overlay of helix 8 from various structures of human SMO (PDB IDs: 5L7D, 6O3C, 6D32), showing that the orientation of the two tryptophan residues into the hydrophobic core of the membrane is conserved (see additional structures in **Fig. S4D**). For consistency with the GST fusions used in capture assays, residue numbering corresponds to mouse SMO.

### A conceptual model for BBSome binding to cargoes and membranes based on mapping of the SMO–BBSome interaction and the cryo-EM structure

To gain insights into how the BBSome enables TZ crossing of its attached cargoes, we sought to model the binding of the BBSome to membranes and cargoes. We started by mapping the interaction of the BBSome with its well-characterized cargo SMO. The BBSome directly recognizes the cytoplasmic tail of SMO that emerges after the seven-transmembrane helix bundle (SMO^Ctail^, aa 543-793; Klink et al., 2017; Seo et al., 2011) and is required for the constitutive removal of SMO from cilia (Eguether et al., 2014; Goetz et al., 2017; Zhang et al., 2011, p. 3). Using *in vitro*-translated (IVT) BBSome subunits, we found that BBS7 was the only subunit detectably captured by SMO^Ctail^ (**Fig. 3A**). BBS7 was also the sole subunit to recognize SSTR3^i3^ (**Fig. S4A**). Truncations of SMO^Ctail^ revealed that the first 19 amino acids of SMO^Ctail^ are necessary and sufficient for binding to BBS7 (**Fig. 3B**). The specificity of BBS7 binding to SMO was retained when BBSome subunits were assayed against the first 19 amino acids of SMO^Ctail^ (**Fig. S4B**). Systematic yeast two-hybrid (YTH) testing using a collection of well-validated constructs (Woodsmith et al., 2017) identified a direct interaction between SMO^Ctail^ and a BBS7 fragment C-terminal to the β-propeller (BBS7[326-672]; BBS7^C^) (**Fig. 3C**). Again, deletion of the first 10 amino acids from SMO^Ctail^ abolished the YTH interaction with BBS7^C^ (**Fig. 3D**). In close agreement with our findings, BBS7 is one of only two BBSome subunits associating with SMO^Ctail^ in co-IP studies and deletion of the first 10 amino acids of SMO^Ctail^ abolishes the interaction with BBS7 (Seo et al., 2011). The congruence of co-IP, YTH and GST/IVT-capture assays strongly supports the conclusion that the first 10 amino acids from SMO^Ctail^ and BBS7 are the major determinants of the SMO–BBSome interaction.

The location of the BBSome-binding determinant on SMO is surprising because the crystal structures of SMO have revealed that the first 10 amino acids of SMO^Ctail^ form a membrane-parallel amphipathic helix termed helix 8 (H8) (Byrne et al., 2016; Deshpande et al., 2019; Huang et al., 2018; Qi et al., 2019; Wang et al., 2014, 2013; Weierstall et al., 2014; Zhang et al., 2017). We thought to determine how the BBSome recognizes SMO^H8^ by mapping the residues of SMO^H8^ required for association with BBS7. A prior study found that the BBSome recognizes motifs consisting of an arginine preceded by an aromatic residue (Klink et al., 2017). One motif in SMO^H8^ is a perfect match (549WR550) and another is a looser candidate (553WCR555). Mutation of both Trp549 and Trp553 from SMO^H8^ completely abolished binding to BBS7, with Trp549 being the major determinant (**Fig. 3E**). Similarly, mutation of Arg550 greatly diminished binding to BBS7. We conclude that each residue within the 549WR550 motif contributes to BBSome binding. The direct binding of BBS7 to Trp549 of SMO was unexpected, because all crystal structures of SMO point to these residues being embedded within the hydrophobic core of the lipid bilayer (**Fig. 3F, S4D**). It follows that SMO^H8^ must be extracted from the membrane for the BBSome to recognize SMO.

Amphipathic helices generally fold upon insertion into the membrane and remain as random coil in solution (Drin and Antonny, 2010; Seelig, 2004). Helix 8 is a near-universal feature of GPCRs (Piscitelli et al., 2015) and a peptide corresponding to helix 8 of rhodopsin adopts a helical conformation when bound to membranes but is a random coil in solution (Krishna et al., 2002). Such membrane requirements for folding of helix 8 are likely generalizable to other GPCRs (Sato et al., 2016). We therefore propose that SMO^H8^ exists in an equilibrium between a folded membrane-embedded and an unfolded state, and that it is the out-of-the-membrane, unfolded state that binds to the BBSome.

Further mapping of the SMO^H8^–BBS7 interaction by GST/IVT-capture assays indicated that BBS7^βprop^ was necessary and sufficient for the interaction with SMO^H8^ (**Fig. 4A**). In YTH, deletion of BBS7^cc^ specifically abolished the SMO^Ctail^–BBS7^C^ interaction (**Fig. 4B**). Thus, while YTH and GST/IVT-capture assays identified different regions of BBS7 recognizing SMO^Ctail^, the regions identified by YTH (BBS7^cc^) and GST/IVT capture (BBS7^βprop^) are adjacent to one another in the BBSome structure. To unify these findings, we propose that an extended SMO^H8^ is recognized by a surface encompassing BBS7^cc^ and BBS7^βprop^.

**Figure 4.**
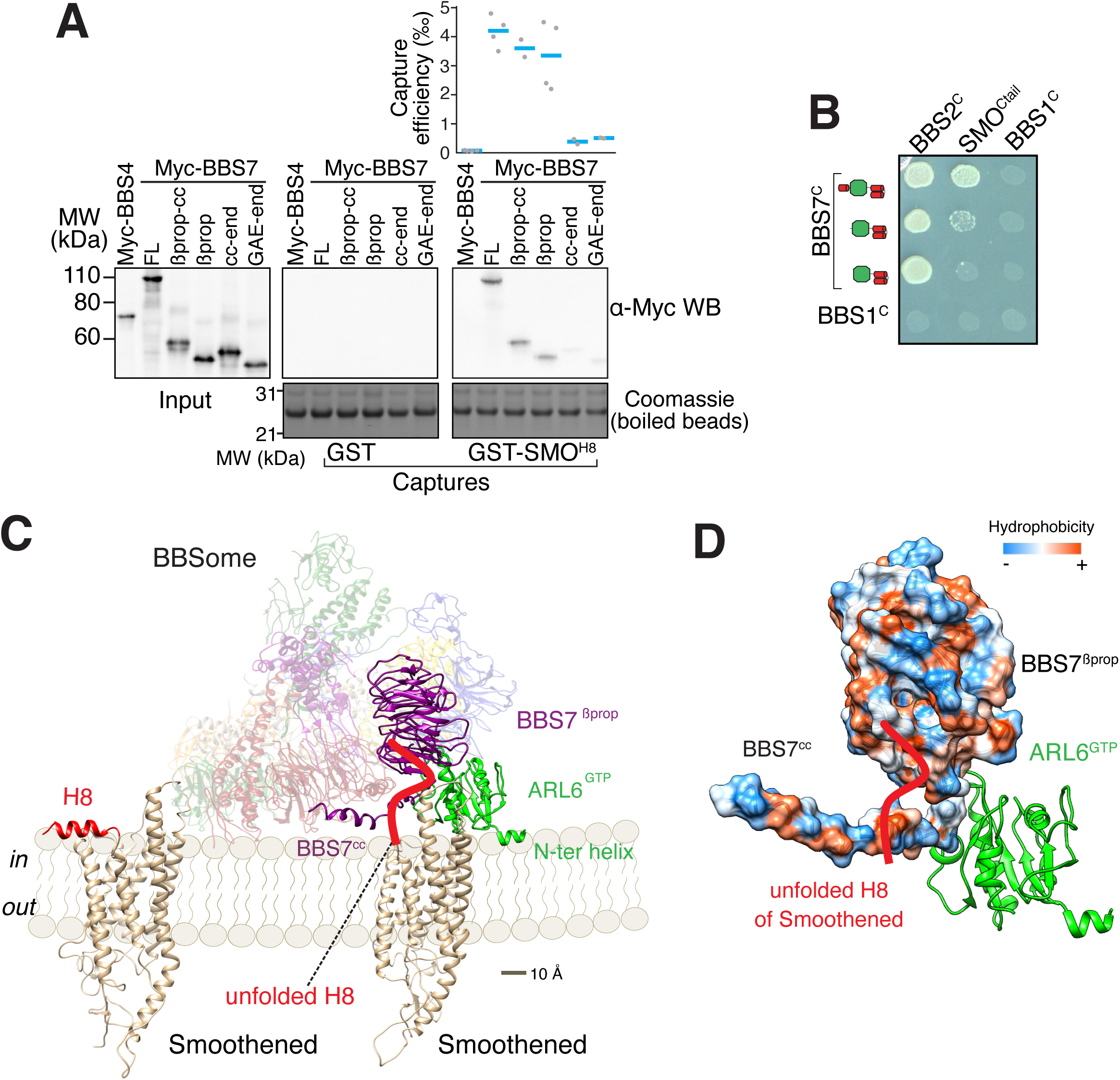
A model for binding of the BBSome to membranes and cargo. **A.** Capture assays of BBS7 find that BBS7^βprop^ engages SMO^H8^. The boundaries of each truncation are βprop, aa 1-332; βprop-cc, aa 1-378; cc-end, aa 326-672; and GAE-end, aa 375-672. Results are presented as in Fig. 3. The sequences of the peptides fused to GST after the protease cleavage site are indicated. **B.** YTH assays show that the deletion of BBS7^cc^ impairs the interaction of BBS7^C^ with SMO^Ctail^ (top row), but not with a C-terminal fragment of BBS2 (BBS2^C^, residues 324-712) (middle rows). BBS1^C^ serves as non-interacting control. Growth controls on diploid-selective medium are shown in **Fig. S5A. C**. Diagram illustrating the proposed interaction of SMO with the membrane-bound BBSome–ARL6^GTP^ complex. For clarity, ARL6^GTP^ and the BBS7 domains involved in SMO binding (BBS7^βprop^ and BBS7^cc^) are shown in solid colors with the remaining subunits shown with reduced opacity. Helix 8 (H8) of Smoothened by itself is folded, whereas it is proposed to become unfolded in the Smoothened– BBSome complex. **D.** Hydrophobicity surface of the BBS7^βprop^ and BBS7^cc^ domains, showing a plausible binding surface for unfolded SMO^H8^ (shown in red). ARL6^GTP^ is shown in ribbon representation.

We conceptualized BBSome binding to membrane and SMO based on our binding studies. Taking into account that Trp549 of SMO is contacted by BBS7^βprop^ and that the first amino acid of SMO after the 7^th^ transmembrane helix is Lys543, BBS7^βprop^ must be within 6 amino acids or ∼21 Å of the membrane. Similarly, BBS7^cc^ must be present within 10 amino acids of the membrane. We note that if SMO^H8^ were to remain helical once extracted from the membrane, BBS7^βprop^ would need to be within 9 Å of the membrane. Because no BBSome–ARL6 orientation can be achieved that brings BBS7^βprop^ within less than 15 Å of the membrane, we conclude that SMO^H8^ must be unfolded to be recognized by the BBSome. Secondly, ARL6^GTP^ anchors the BBSome to the membrane. Because ARF family GTPases bind lipid bilayers through their amphipathic N-terminal helix inserted in a membrane-parallel orientation within the lipid-headgroup layer, the starting point of the core GTPase domain of ARL6 at Ser15 informs the anchoring of the BBSome–ARL6^GTP^ complex on membranes. In the resulting conceptual model of the membrane-associated BBSome–ARL6^GTP^ complex bound to SMO (**Fig. 4C**), the orientation with respect to the membrane of ARL6^GTP^ in complex with the BBSome is similar to that of other Arf-like GTPases in complex with coat adaptor complexes (**Fig. S5B**) (Cherfils, 2014). The BBSome–ARL6^GTP^ complex displays a convex membrane-facing surface, defined by the N terminus of ARL6^GTP^ and parts of BBS2^pf^, BBS7^βprop^ and BBS9^βprop^, that espouses the contour of the ciliary membrane modeled as a 250-nm cylinder (**Fig. 4C**). A convex membrane-binding surface in the Golgin^GRIP^–ARL1^GTP^ or MKLP1– ARF6^GTP^ complexes similarly allows these complexes to associate with concave surfaces (Makyio et al., 2012; Panic et al., 2003). Importantly, a hydrophobic cluster can be traced through the surfaces of BBS7^cc^ and BBS7^βprop^ to make a strong candidate for the critical Trp residues in SMO^H8^ (**Fig. 4D**).

### Molecular interactions of the BBSome with membranes and the IFT-B complex

Liposome-recruitment assays with pure BBSome and ARL6^GTP^ have shown that the BBSome recognizes lipid headgroups, in particular the phosphoinositide PI(3,4)P_2_ (Jin et al., 2010). Pleckstrin Homology (PH) domains are prototypical PIP-recognition modules and PIP-overlay assays suggested that BBS5^PH1^ might directly recognize PIPs (Nachury et al., 2007), although it has been noted that PIP-overlay assays can report spurious interactions (Yu et al., 2004). We sought to determine whether the PH domains of BBS5 can bind to lipid headgroups in our model of the membrane- and cargo-bound BBSome. The canonical PIP-binding motif Kx_n_[K/R]xR is present in the β1-β2 loop of nearly all PH domains that bind PIPs (Isakoff et al., 1998; Vonkova et al., 2015). BBS5^PH1^ contains a perfect match to the PIP-binding motif (K41xxxxxR47xR49) but no such motif is found in BBS5^PH2^ (**Fig. 5A**). Consistent with the absence of a PIP-binding motif in BBS5^PH2^, lipid binding is blocked by the edge of a blade from BBS9^βprop^ (**Fig. 5B**). When the canonical PIP-binding site was mapped to the structure of BBS5^PH1^, the lipid-binding site was occluded by a loop connecting BBS7^βprop^ to BBS9^cc^ (**Fig. 5B**). Modeling 9 distinct PH domains co-crystallized with PIP headgroups onto BBS5^PH1^ showed limited variance in the lipid orientation (**Fig. 5C**). In summary, the PH domains of BBS5 are unable to recognize PIP through their canonical sites.

**Figure 5.**
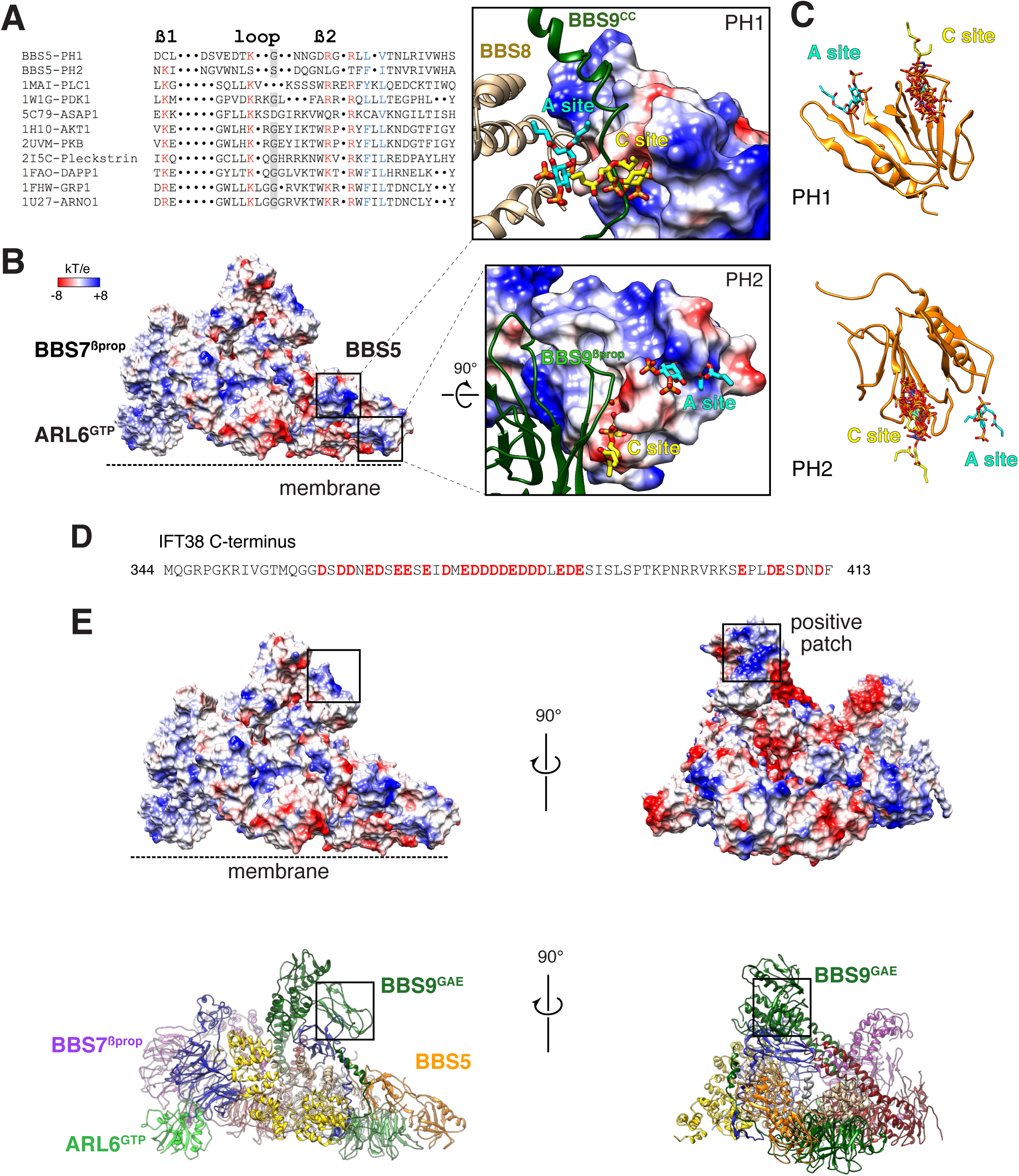
Mapping of potential molecular interactions of the BBSome with lipids and IFT onto the structure. **A.** Sequence alignment of the β1-loop-β2 region in structurally characterized PH domains. Conserved residues are highlighted; grey background shading, glycine; red font, positively charged residues; blue font, hydrophobic residues. **B.** Electrostatic surface of the membrane-bound BBSome–ARL6^GTP^ complex, and close-up views of the BBS5 pleckstrin homology (PH) domains. For the PH1 domain, the canonical (C) and atypical (A) sites for lipid binding are occluded by BBS9 (dark green ribbon) and BBS8 (gold ribbon), respectively. For the PH2 domain, the C site is blocked by BBS9, but the A site is accessible. The lipids at the A and C sites, shown as yellow and cyan sticks, respectively, are diC4-PtdIns(4,5)P2 and are modeled based on the structural alignment of the BBS5 PH domains with the lipid-bound ASAP1 PH domain (PDB ID: 5C79) **C.** Structural overlay of the lipids bound to the PH domains listed in the above sequence alignment (PDB IDs: 1MAI, 1W1G, 5C79, 1H10, 2UVM, 2I5C, 1FAO, 1FHW, 1U27), showing the consistency of the lipid position for both PH domains. The lipids are shown as stick models and are overlaid on the PH1 and PH2 domains of BBS5 (orange ribbon). **D.** Sequence of the region of IFT38 that interacts with the BBSome. See **Fig. S6D** for domain organization of IFT38. Negatively charged residues are bold and red. **E.** Top: Electrostatic surface of the ARL6^GTP^ -bound BBSome showing a patch of positive charges on a region of BBS9 that interacts with BBS1 and BBS2 and is a candidate for binding IFT38^C^. Bottom: corresponding orientations in ribbon diagram representations.

More recently, the crystal structure of the PH domain of ASAP1 has provided singular evidence for an atypical PIP-binding (A) site (Jian et al., 2015). In BBS5^PH1^, lipid binding to the predicted A site extensively clashes with BBS8^TPR8-9^ (**Fig. 5B**). In BBS5^PH2^, lipid binding to the putative A site would cause no steric clash (**Fig. 5B**). However, considering that the distance between the membrane and the A site of BBS5^PH2^ exceeds 1 nm and considering the limited evidence for the existence of A sites in PH domains, it is very unlikely that the A site of BBS5^PH2^ participates in lipid binding of the BBSome.

Besides the PH domains of BBS5, we inspected the membrane-facing surface of the BBSome for positively charged surfaces (**Fig. S6A**). Interestingly, the surface of the ARL6^GTP^-bound BBSome exhibited considerably fewer negative charges facing the membrane than the surface of BBSome alone (compare **Fig. S6A** and **S6B**). While some negatively charged surfaces facing the membrane remained, the surfaces closest to the membrane are generally positively charged in the BBSome–ARL6^GTP^ complex structure.

We next considered binding of the BBSome to IFT. IFT38/CLUAP1 is the only IFT-B subunit to consistently interact with the BBSome in systematic affinity purification studies (Boldt et al., 2016) and a recent study found that the C terminus of IFT38 interacts with the BBSome in visual immunoprecipitation (VIP) assays (Nozaki et al., 2019). Using GST-capture assays with pure BBSome, we confirmed that IFT38 directly interacts with the BBSome and that the C-terminal domain of IFT38 is necessary and sufficient for this interaction (**Fig. S6C** and **D**). The C-terminal tail of IFT38 (aa 329-413) is predicted to be unstructured and is remarkably acidic in that 30 of its 85 amino acids are either Glu or Asp, giving it a theoretical pI of 4.03. As VIP assays identified BBS9 as the major binding subunit of IFT38^C^, with contributions from BBS2 and BBS1, we reasoned that a BBS9 domain in close proximity to BBS2 and BBS1 should be responsible for IFT38^C^ binding. The C terminus of BBS9 (GAE, pf, hp, CtH) sits atop BBS1^GAE^ and connects to BBS2^hp^ and an extended positive patch is found in BBS9^GAE^ (**Fig. 5E**). This positive patch is therefore a strong candidate for the IFT38^C^ interaction and its orientation away from the membrane makes it well-positioned to interact with IFT trains associated with the axoneme.

### A hypothesis for BBSome-mediated passage of GPCRs across the transition zone

Our combination of binding data and structural studies leads to the conclusion that SMO^H8^ must be extracted from its location in the plane of the inner leaflet and unfolded in order for SMO to be recognized by the BBSome (**Fig. 4C**). We considered how switching this region of SMO from membrane-embedded amphipathic helix to random coil may underlie selective crossing of the TZ. It is now established that amphipathic helices require headgroup packing defects to insert themselves into the cytoplasmic leaflet of cellular membranes (Bhatia et al., 2010; Drin and Antonny, 2010; Hatzakis et al., 2009). Lipid composition and membrane curvature are the two factors that influence the density of headgroup-packing defects. As membranes curve toward the cytoplasm (positive curvature), the distance between neighboring headgroups in the cytoplasmic leaflet increases and interfacial membrane voids begin to appear. Amphipathic helices therefore preferentially insert into positively curved membranes and this forms the basis for the recruitment of many proteins to nascent endocytic buds (Larsen et al., 2015; Miller et al., 2015). Meanwhile, ultrastructural views of the TZ from organisms as diverse as *Paramecium*, *Chlamydomonas*, *Trypanosome*, *Drosophila* and mammals show that the membrane of the TZ holds a rigidly maintained negative curvature (**Fig. 6A**) (Fisch and Dupuis Williams, 2011). The numerous contacts between TZ proteins and TZ membrane are likely to brace the TZ membrane and force a rigid negative curvature (Garcia-Gonzalo and Reiter, 2017). In contrast, the membrane of the ciliary shaft appears considerably more tolerant to local deformations (**Fig. 6A**).

**Figure 6.**
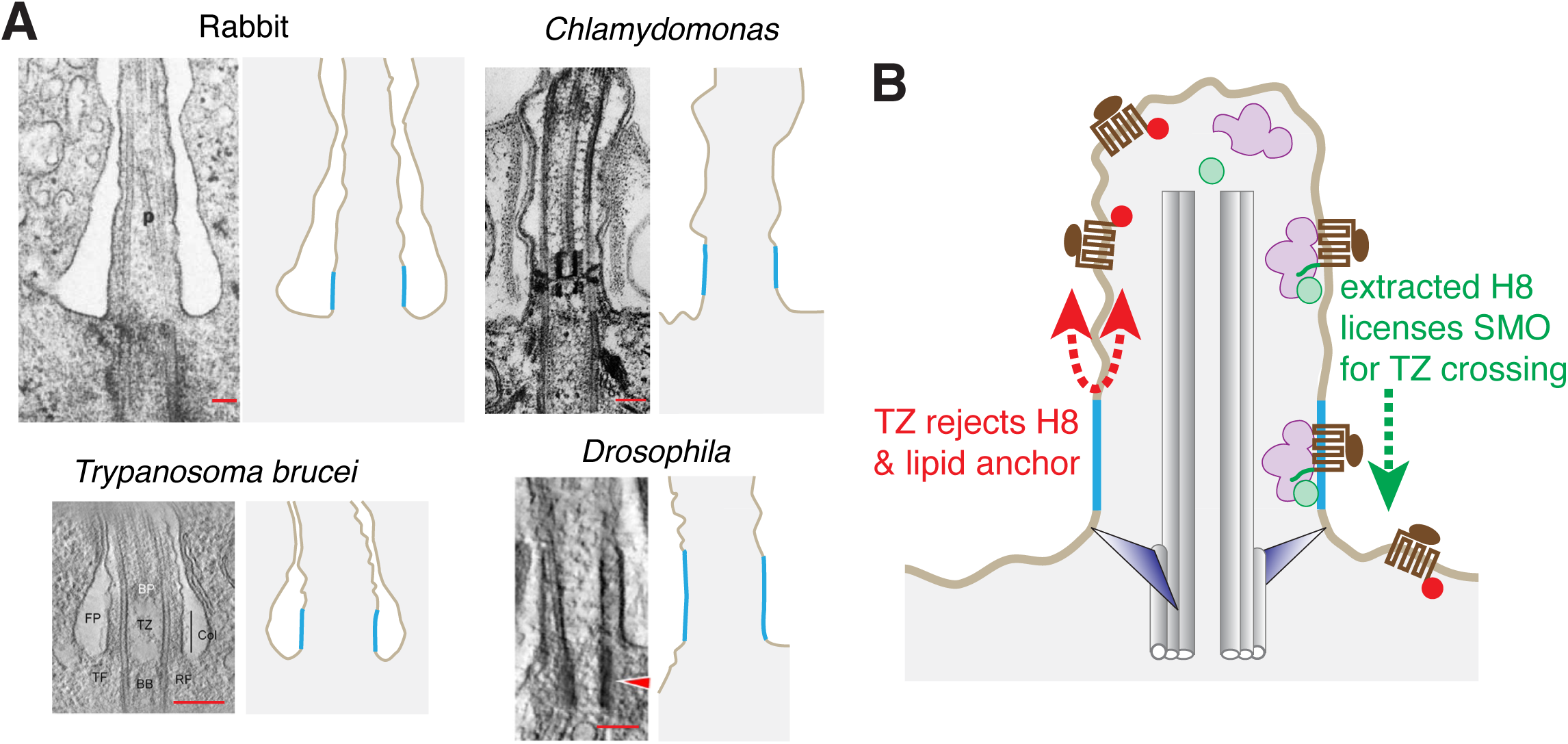
A model for selective crossing of the transition zone (TZ) based on the rejection of inner leaflet insertions by the transition zone membrane. **A.** Thin-section electron micrographs of the TZ from evolutionarily diverse organisms (left panels) and schematic drawings (right panels) illustrate the rigid membrane at the TZ (blue) as compared to the irregular ciliary shaft membrane (brown). Images are reprinted from (Gallagher, 1980; Jana et al., 2018; Ringo, 1967; Trépout et al., 2018). Scale bar: 200 nm. **B.** A model for selective passage through the TZ. The amphipathic helix 8 of ciliary GPCRs is tolerated in the flexible membrane of the ciliary shaft but not in the rigid outwardly curved membrane of the TZ. By extracting helix 8 out of the inner leaflet, the BBSome licenses GPCRs to diffuse through the TZ.

Secondly, headgroup packing defects are favored by some lipid compositions – e.g., lipids with small headgroups – but are largely depleted in liquid-ordered phases (Drin and Antonny, 2010; Larsen et al., 2015). Congruently, ARF1 and its N-terminal amphipathic helix are excluded from liquid-ordered domains in artificial membranes (Manneville et al., 2008). The findings of a condensed lipid phase at the base of the cilium (Schou et al., 2017; Vieira et al., 2006) and the observation that the TZ membrane is the most resistant membrane to detergent extraction in *Chlamydomonas* (Kamiya and Witman, 1984) indicate that the TZ membrane has liquid-ordered properties and will thus exclude amphipathic helices. Together, compositional and ultrastructural features of the TZ suggest that a negatively curved and tightly packed membrane at the TZ acts as a filter against amphipathic helices inserted into the cytoplasmic leaflet. By increasing the energetic cost of inserting amphipathic helices into the inner leaflet, the geometry and composition of the TZ membrane will impede the diffusion of GPCRs containing a membrane-embedded helix 8 through the TZ (**Fig. 6B**). As the membrane of the ciliary shaft appears considerably more tolerant to local deformations (**Fig. 6A**), it will tolerate insertion of SMO^H8^ in the inner leaflet. In this context, SMO^H8^ can be viewed as a toggle switching between a membrane-embedded orientation that is rejected from diffusing into the TZ and an extracted, BBSome-associated conformation competent to cross the TZ (**Fig. 6B**).

### Increasing the depth of insertion of the ARL6 amphipathic helix blocks BBSome-mediated exit

In contrast to SMO^H8^, the amphipathic helix at the N terminus of ARL6 contains no bulky hydrophobic groups (**Fig. 7A**). It is thus conceivable that the energetic cost of inserting the amphipathic helix of ARL6 into the membrane of the TZ is tolerated but that inserting SMO^H8^ in the TZ membrane is prohibitive. To test the hypothesis that amphipathic helices deeply inserted into the inner leaflet see the TZ as a roadblock, we swapped the N-terminal 13 amino acids of ARL6 for the N-terminal 15 amino acids of ARF1 (**Fig. 7B**). While the amphipathic helix of ARL6 contains no aromatic residues, the amphipathic helix of ARF1 contains three phenylalanine residues (**Fig. 7A**). Molecular dynamics simulations have shown that the amphipathic helix of ARF1 reaches halfway into the inner leaflet while milder amphipathic helices surf on the surface of the membrane (Li, 2018).

**Figure 7.**
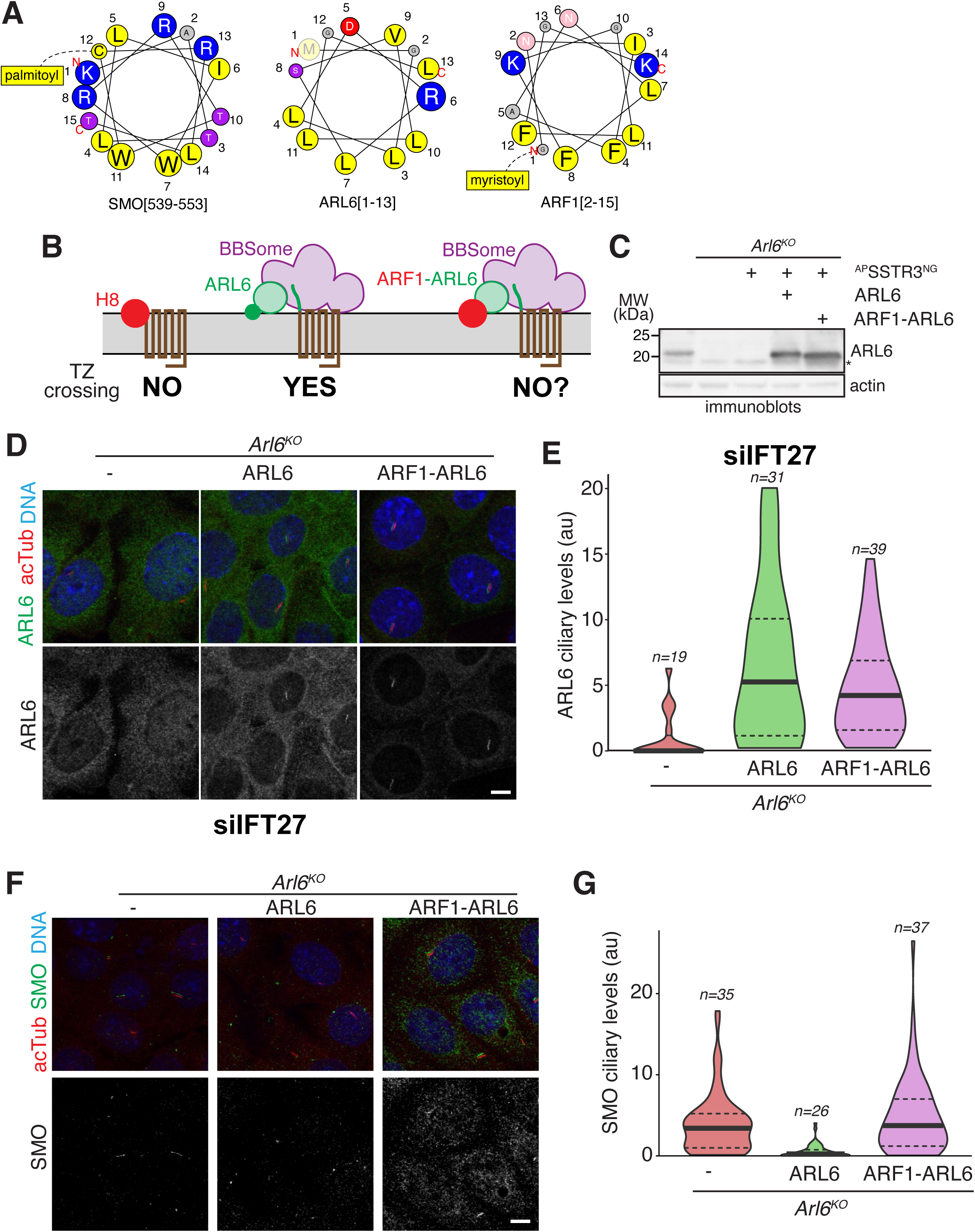
A chimera between the amphipathic helix of ARF1 and the GTPase domain of ARL6 localizes to cilia but fails to support SMO exit from cilia. **A**. Helical wheel representations of the amphipathic helices of ARL6, SMO and ARF1. The size of the circles is proportional to the relative size of the residues. Also shown are the palmitoyl group attached to Cys554 of SMO and the myristoyl group attached to Gly1 of ARF1. **B**. Diagram of the predictions made by the helix rejection model. Amphipathic helices that penetrate deeply into the aliphatic core of the membrane (SMO^H8^, ARF1^N^; large red solid circles) are rejected from the inner leaflet of the transition zone while weakly amphipathic helices that do not penetrate deeply into the membrane (ARL6^N^; small green solid circle) are tolerated by the transition zone. **C-E.** Stable and clonal cell lines were generated in IMCD3-FlpIn *Arl6^-/-^* cells by introducing ARL6 or a chimera in which the amphipathic helix of ARL6 is replaced by that of ARF1 (ARF1-ARL6). **C.** Expression levels of the ARL6 variants were assessed by immunoblotting. Actin was used as a loading control. In the rescued cell lines, ARL6 is expressed at 2.6-fold the endogenous level, while ARF1-ARL6 is expressed at 3.2-fold the endogenous level. **D.** Cells stained for ARL6, acetylated tubulin and DNA show that ARF1-ARL6 localizes to cilia. Cells were depleted of IFT27 by siRNA to bring the levels of ARL6 above the detection threshold of immunostaining. See **Fig. S7A-B** for control-depleted cells. Scale bar: 10 μm. **E.** The fluorescence intensity of the ARL6 channel in the cilium was measured in each cell line and the data are represented in a violin plot. The thick bar indicates the median and the dotted lines the first and third quartiles. **F.** Cells stained for SMO, acetylated tubulin and DNA show that ARF1-ARL6 fails to support SMO exit from cilia. Cells were not treated with Hedgehog pathway agonist. See **Fig. S7C-D** showing the same cell lines treated with the Smoothened agonist (SAG). Scale bar: 10 μm. **G.** The fluorescence intensity of the SMO channel in the cilium was measured in each cell line and the data are represented in a violin plot.

Stable clonal cell lines expressing ARL6 or the ARF1-ARL6 chimera were generated in murine IMCD3 kidney cells in which *Arl6* had been deleted. Care was exercised to limit expression levels by leveraging single-site genomic integration and a weak promoter, resulting in levels of reintroduced ARL6 proteins that were within 3-fold of endogenous ARL6 levels in wild-type IMCD3 cells (**Fig. 7C**). To determine the functionality of ARF1-ARL6, we assessed its ciliary localization and found that it localized to cilia to the same extent as the reintroduced ARL6 (**Fig. 7D** and **E**).

Next, we assayed the constitutive removal of SMO from cilia by the BBSome–ARL6^GTP^ complex. Immunostaining for endogenous SMO confirmed that SMO accumulates in cilia of unstimulated *Arl6^-/-^* cells and that reintroduced ARL6 brings down the ciliary SMO levels below the detection limit of immunostaining (**Fig. 7F** and **G**). In contrast, ARF1-ARL6 failed to rescue the ciliary removal of SMO in unstimulated cells as SMO accumulated in cilia to the same extend in ARF1-ARL6 cells as in *Arl6^-/-^* cells (**Fig. 7F** and **G**). Meanwhile, SMO accumulated in cilia in the presence of SMO agonist to the same extent in all cell lines tested (**Fig. S7C** and **D**). We conclude that ARF1-ARL6 is unable to support BBSome-dependent removal of SMO from cilia, despite the ability of ARF1-ARL6 to localize to cilia.

We next investigated whether the amphipathic helix exclusion model may be generalizable to other GPCRs besides SMO. Searching for the BBSome-binding motif [W/F/Y][K/R] (Klink et al., 2017) within H8 of the 26 known ciliary GPCRs revealed that 23 of them contained a BBSome-binding motif in their helix 8 (**Fig. 8A**). Considering that aromatic residues will point towards the core of the membrane in these H8, the broad distribution of BBSome-binding motifs suggests that the BBSome may bind to extracted H8 in nearly all ciliary GPCRs. As signal-dependent exit of GPCRs has been observed in nearly every GPCR tested (Nachury, 2018), it is tempting to generalize the model of H8 extraction as a licensing mechanism for TZ crossing. We zeroed in on the somatostatin receptor 3 (SSTR3), given its well-characterized, BBSome-dependent exit from cilia (Green et al., 2015; Ye et al., 2018). SSTR3 was introduced into the IMCD3 cell lines together with the ARL6 proteins and its exit was monitored thanks to a fluorescent protein (mNeonGreen) fused to its intracellular C-terminal tail. As in the case of SMO, SSTR3 exit failed in *Arl6^-/-^* and ARF1-ARL6 cells but was rescued to normal kinetics in ARL6 cells (**Fig. 8B**). We conclude that the very shallow insertion of the ARL6 amphipathic helix into the inner leaflet is required for ARL6 to support BBSome-dependent exit from cilia. Together, these results support the model that extraction of GPCR H8 from the inner leaflet is a required step in BBSome-mediated exit from cilia.

**Figure 8.**
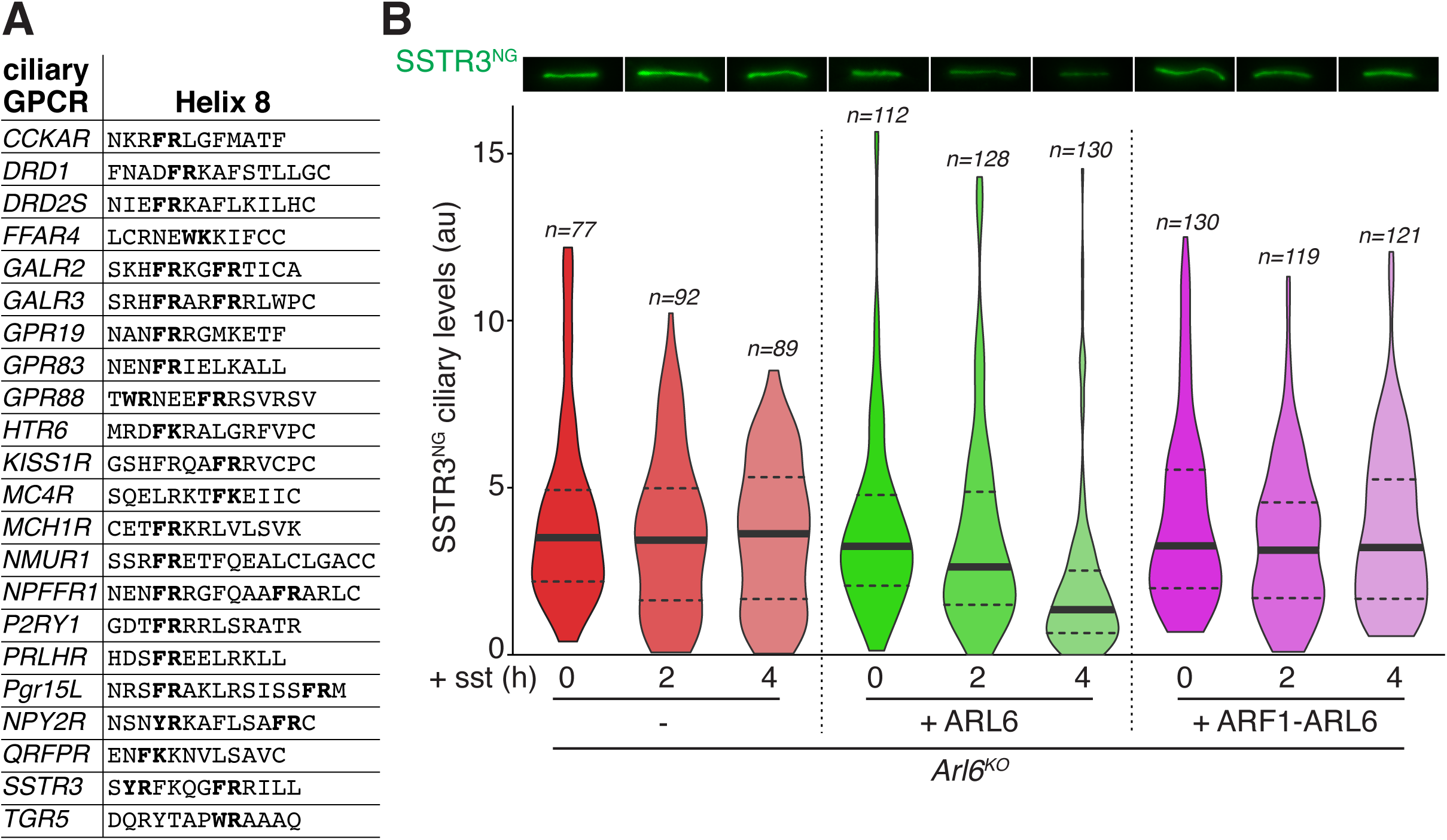
A chimera between the amphipathic helix of ARF1 and the GTPase domain of ARL6 fails to support SSTR3 exit. **A.** Sequence analysis finds a BBSome-binding motif ([W/F/Y]R) within helix 8 in 20 of the 26 GPCRs known to localize to cilia. **B.** SSTR3 exit assays in the ARL6 rescue lines. Cells were treated with the SSTR3 agonist somatostatin for the indicated time points, fixed and the ciliary fluorescence of NeonGreen-tagged SSTR3 (SSTR3^NG^) was measured. Micrographs of representative individual cilia are shown on top. The ciliary fluorescence intensity of SSTR3^NG^ was measured in each cell line and the data are represented as violin plots.

## DISCUSSION

### Biochemical properties of the open conformation

The specific interaction that is gated by the activating conformational change remains unknown. Mapping of characterized interactions on the BBSome surface suggests that the known BBSome interactors will bind equally well to the open and closed forms.

First, IFT38 interacts with the BBSome in systematic immuno-precipitation/mass spectrometry studies, in VIP assays, and in GST capture of pure BBSome. The candidate interacting region is diametrically opposite from the ARL6-binding region and does not undergo any measurable change upon conformational opening.

Second, BBS7^βprop^ and BBS7^cc^ appear equally accessible in the closed and open conformation (**Fig. 1A, 2A and Video 1**), suggesting that the conformational change is unlikely to directly increase the affinity of BBSome for its cargo SMO. This would contrasts with findings of increased affinity of COPI, AP1 and AP2 for their cargoes upon conformational changes induced by membrane recruitment (Dodonova et al., 2017; Jackson et al., 2010; Ren et al., 2013).

We conclude that the biochemical interaction that is modulated by ARL6^GTP^ binding is likely distinct from cargo or IFT binding. More sensitive biochemical assays or the discovery of novel interactions may be necessary to decipher the interactions that are modulated by conformational opening of the BBSome.

Surprisingly, deletion of the BBSome-binding domain from IFT38 in cells did not grossly alter BBSome distribution in cilia or affect the ability of the BBSome to constitutively remove SMO from cilia, but it did interfere with GPR161 exit (Nozaki et al., 2019). These results suggest that an IFT38 subcomplex distinct from the holo-IFT-B complex may assist the BBSome with specific duties. It is thus conceivable that the interaction of the BBSome with IFT-B is gated by ARL6^GTP^, in line with an increase in retrograde train size that depends on ARL6.

### Generalization of the amphipathic helix exclusion model

Besides their depth of insertion into the membrane, the amphipathic helices of SMO and ARL6 differ in their acylation. While ARL6 is not N-myristoylated (Gillingham and Munro, 2007; Wiens et al., 2010), SMO^H8^ is predicted to be capped by a palmitoylated cysteine (Cys554), similar to helix 8 in class A GPCRs, which are nearly always palmitoylated near their C termini (Piscitelli et al., 2015). More generally, a striking feature common to nearly all BBSome cargoes is the presence of palmitoyl and/or myristoyl anchors (Liu and Lechtreck, 2018). Importantly, headgroup packing defects are also required to accommodate the insertion of lipid anchors attached to a polypeptide chain (Bhatia et al., 2010; Hatzakis et al., 2009; Larsen et al., 2015). The proposed paucity of headgroup packing defects in the cytoplasmic leaflet of the TZ membrane would thus hinder the passage of proteins with attached lipid anchors through the TZ.

The BBSome must shelter a considerable hydrophobic surface when it extracts SMO^H8^ out of the membrane: first, the two tryptophan residues in SMO^H8^ that reside in the hydrophobic core of the lipid bilayer directly bind the BBSome (**Fig. 6** and Klink et al., 2017); second, the palmitoylated Cys554 will find itself outside of the hydrophobic core of the membrane when SMO^H8^ is bound to the BBSome. A cavity that shelters large hydrophobic residues and lipid anchors to hold helix 8 away from the membrane and enable TZ crossing may exist in the BBSome.

## ACKNOWLEDGMENTS

We thank Mark Ebrahim and Johanna Sotiris for help with grid screening and data collection at the Evelyn Gruss Lipper Cryo-Electron Microscopy Resource Center at The Rockefeller University. This work was funded by NIGMS (R01-GM089933, M.V.N.) and Research to Prevent Blindness (Stein Innovator Award A131667, M.V.N). This work was made possible, in part, by NEI (EY002162 - Core Grant for Vision Research, M.V.N.), RPB (Unrestricted Grant, M.V.N.) and the Austrian Science Fund (FWF P30162, U.S.). Molecular graphics and analyses were performed with the UCSF Chimera package. Chimera is developed by the RBVI at UCSF (supported by NIGMS P41-GM103311).

The authors declare no competing interests.

---

**Supplement Figure 1.**
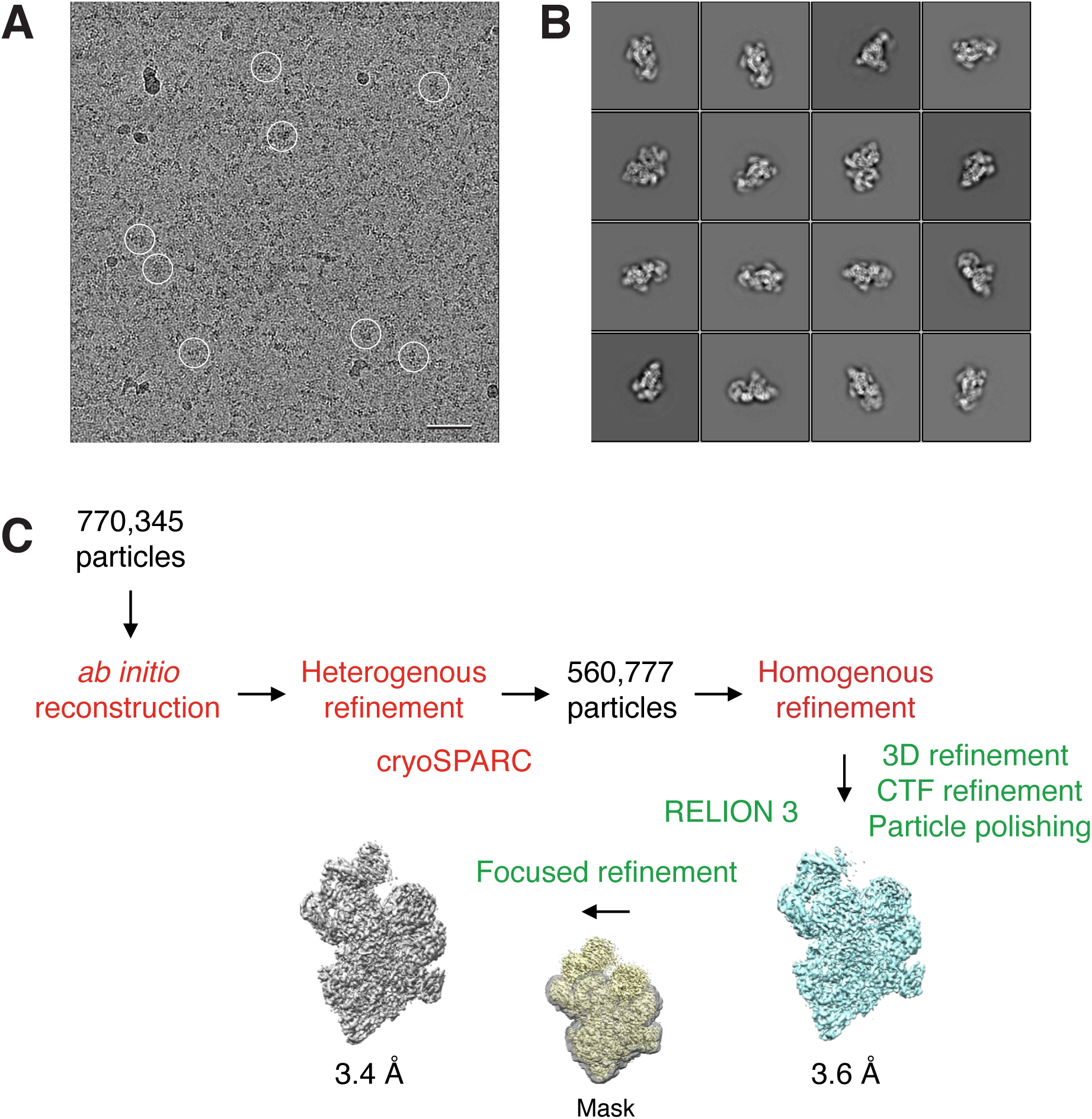
Cryo-EM analysis of the BBSome alone. **A.** Typical cryo-EM image of vitrified BBSome. Some particles are circled. Scale bar: 50 nm. **B.** Selected 2D class averages obtained with RELION 3.0. Side length of individual averages: 41.6 nm. **C.** Image-processing workflow for 3D reconstruction and refinement in cryoSPARC and RELION 3.0. See Methods section for details.

**Supplement Figure 2.**
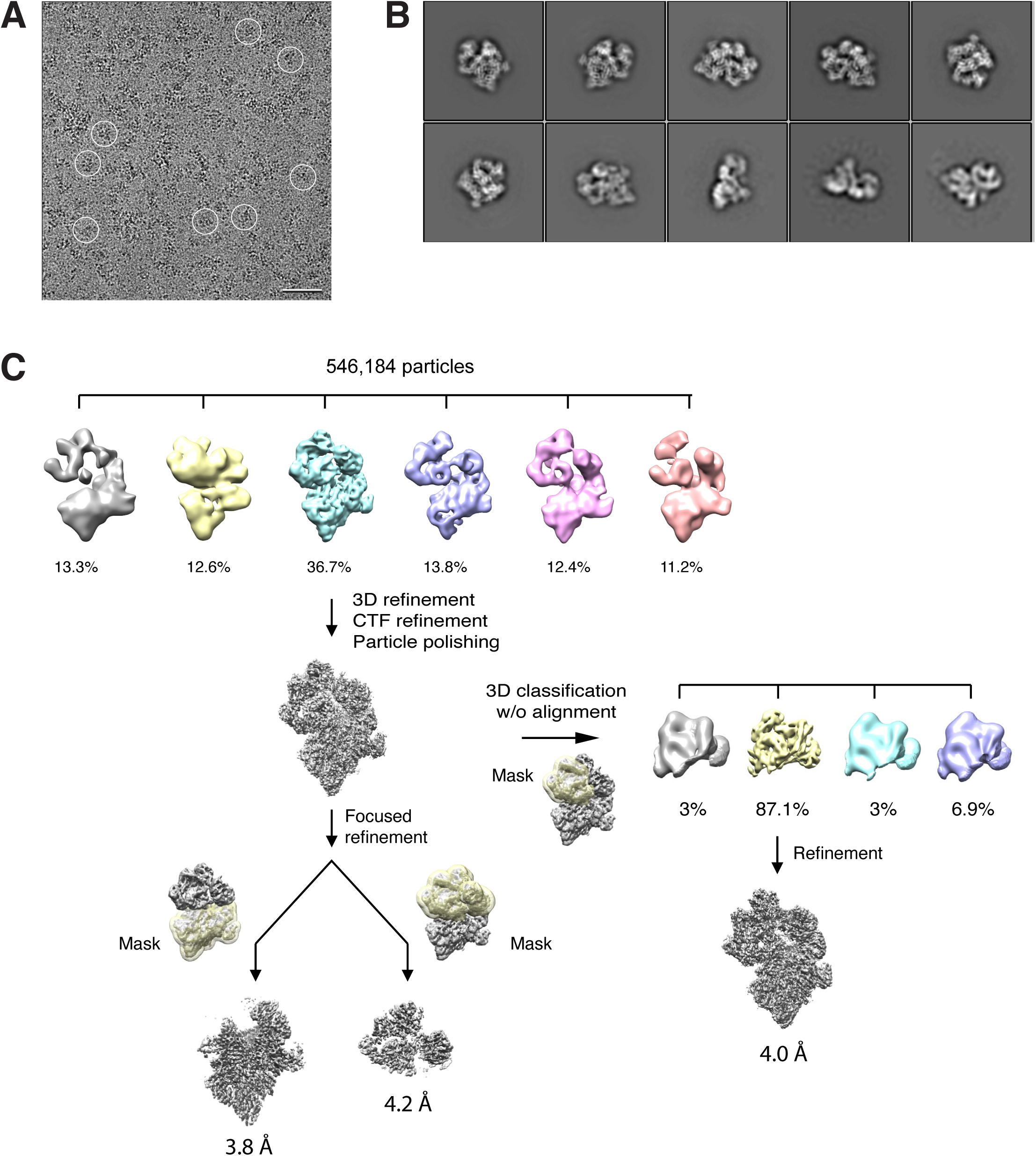
Cryo-EM analysis of the BBSome–ARL6^GTP^ complex. **A.** Typical cryo-EM image of vitrified BBSome–ARL6^GTP^ complex. Some particles are circled. Scale bar: 50 nm. **B.** Selected 2D class averages obtained with RELION 3.0. Side length of individual averages: 45 nm. **C.** Image-processing workflow for 3D classifications and refinement in RELION 3.0 that yielded the three density maps discussed in the main text. See Methods for details.

**Supplement Figure 3.**
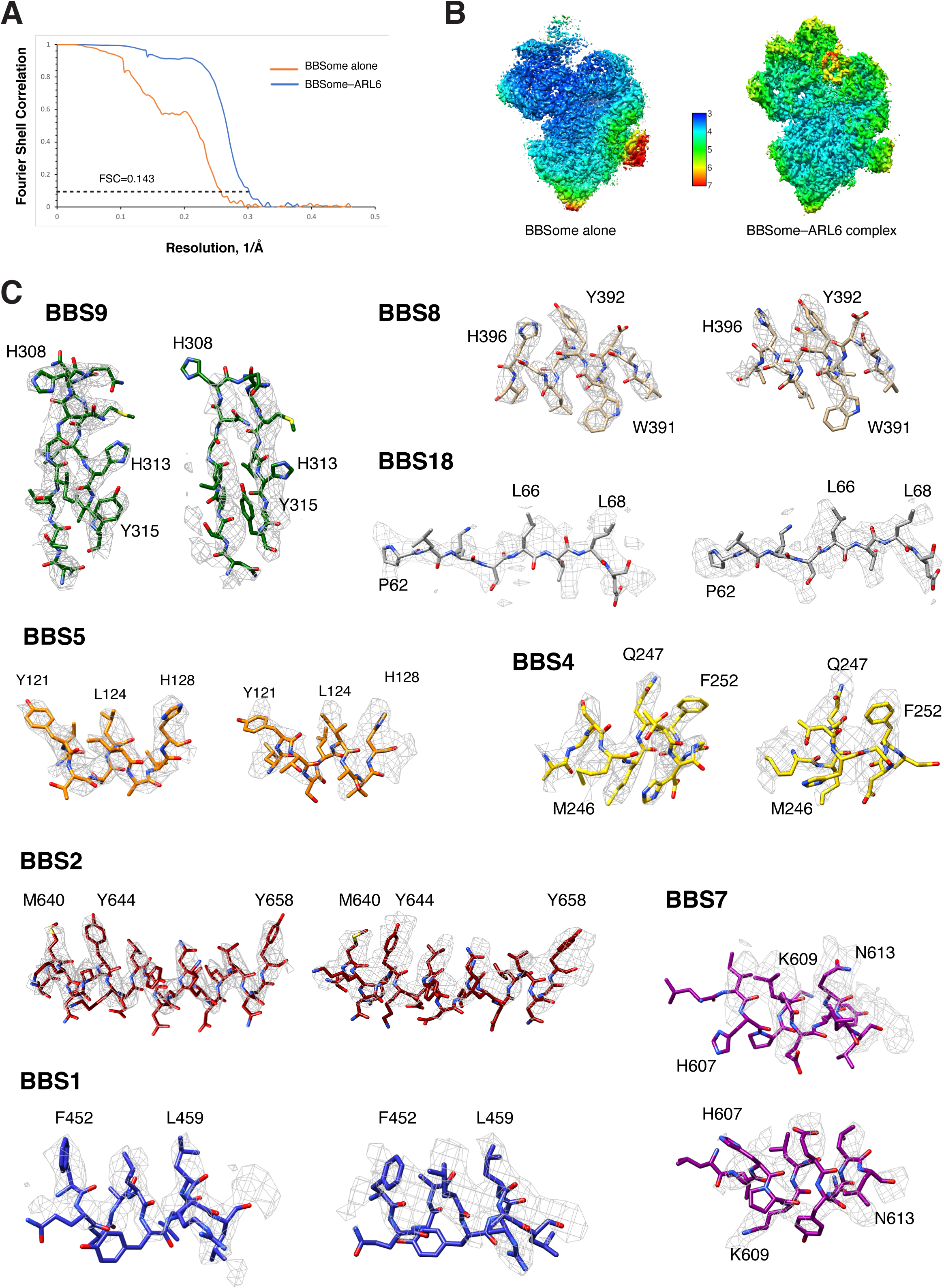
Quality assessment of the BBSome density maps. **A.** Fourier shell correlation curves calculated between independently refined half maps for the density maps of BBSome alone and the BBSome–ARL6^GTP^ complex. **B.** Local resolution for the density maps of BBSome alone and the BBSome–ARL6^GTP^ complex as determined by using the ResMap algorithm included in RELION. **C.** Representative cryo-EM densities for the maps of BBSome alone (left panels) and the BBSome–ARL6^GTP^ complex (right panels).

**Supplement Figure 4.**
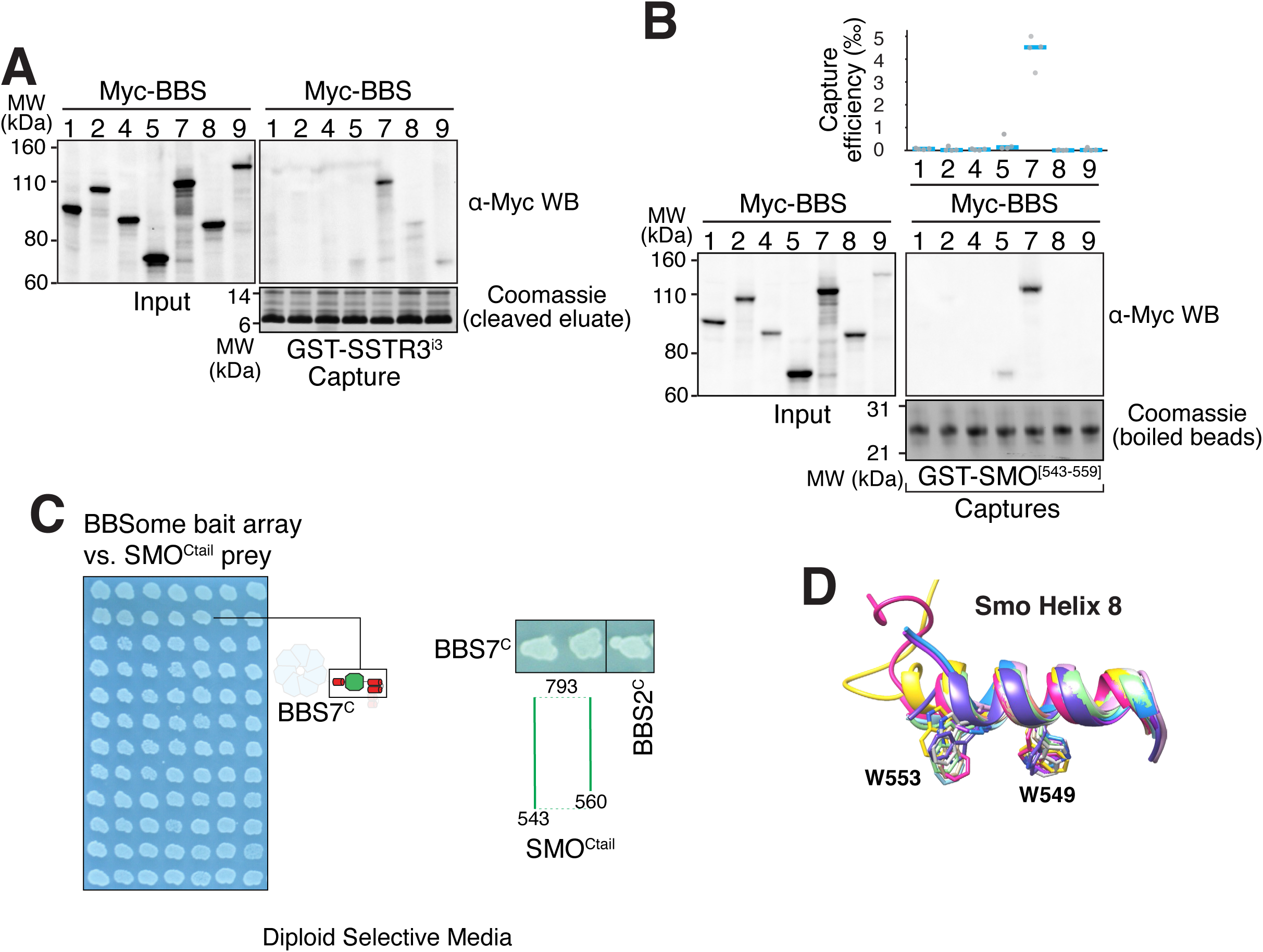
Addition to Figure 3. **A.** Capture of individual BBSome subunits with GST-SSTR3^i3^ identifies BBS7 as the SSTR3^i3^- binding subunit. **B.** Capture of individual BBSome subunits with GST fused to the first 19 amino acids of SMO^Ctail^ (aa 543-559) identifies BBS7 as the SMO-binding subunit. **C**. Controls for diploid growth of the yeast array on medium lacking tryptophan and leucine tested in Fig. 3C-D. **D.** Structural overlay of helix 8 from different human Smoothened proteins (PDB IDs: 4JKV, 4N4W, 4O9R, 4QIM, 4QIN, 5L7D, 5V56, 5V57, 6D32, 6D35, 6OT0). The side chains of the two critical tryptophan residues are shown as sticks. The numbering of the residues is based on mouse Smoothened.

**Supplement Figure 5.**
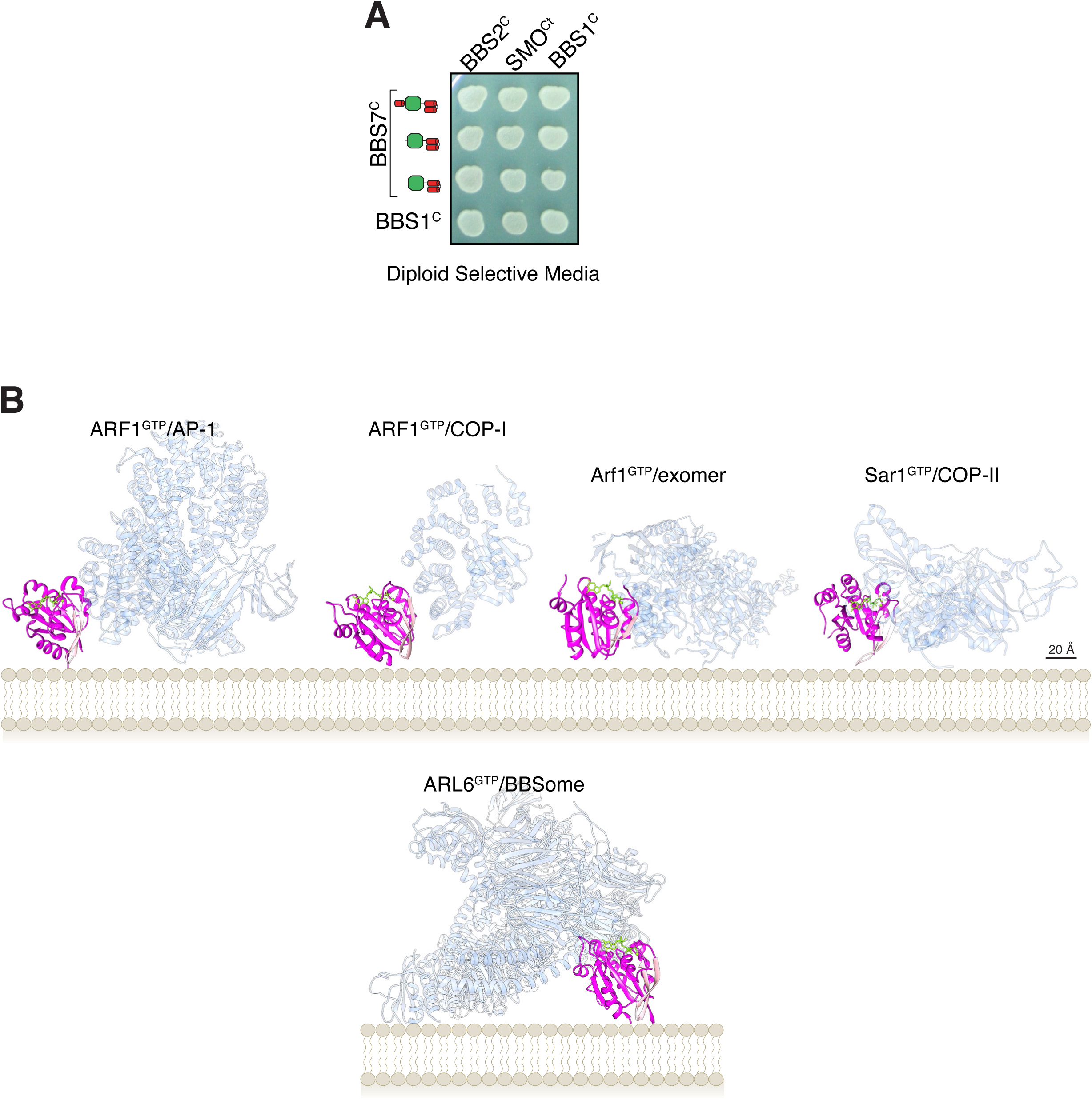
Addition to Figure 4. **A**. Controls for diploid growth of the yeast array on medium lacking tryptophan and leucine tested in Fig. 4B. **B**. Orientations of Arf-like GTPases in complex with coat adaptor complexes from crystal structures (top row) and in our model of the BBSome bound to membranes and cargo. The coat complexes are transparent blue, the GTPases are magenta with their interswitch hairpin in pink, the nucleotide and Mg^2+^ ion are chartreuse. The PDB IDs are: 4HMY for the complex of ARF1^GTP^ with AP1 (Ren et al., 2013), 3TJZ for the complex of ARF1^GTP^ with γζ- COP (Yu et al., 2012), 4Q66 for the complex of Arf1^GTP^ with exomer (Bch1/Chs5) (Paczkowski and Fromme, 2014), and 2QTV for Sar1 bound to Sec23/Sec31 (Bi et al., 2007).

**Supplement Figure 6.**
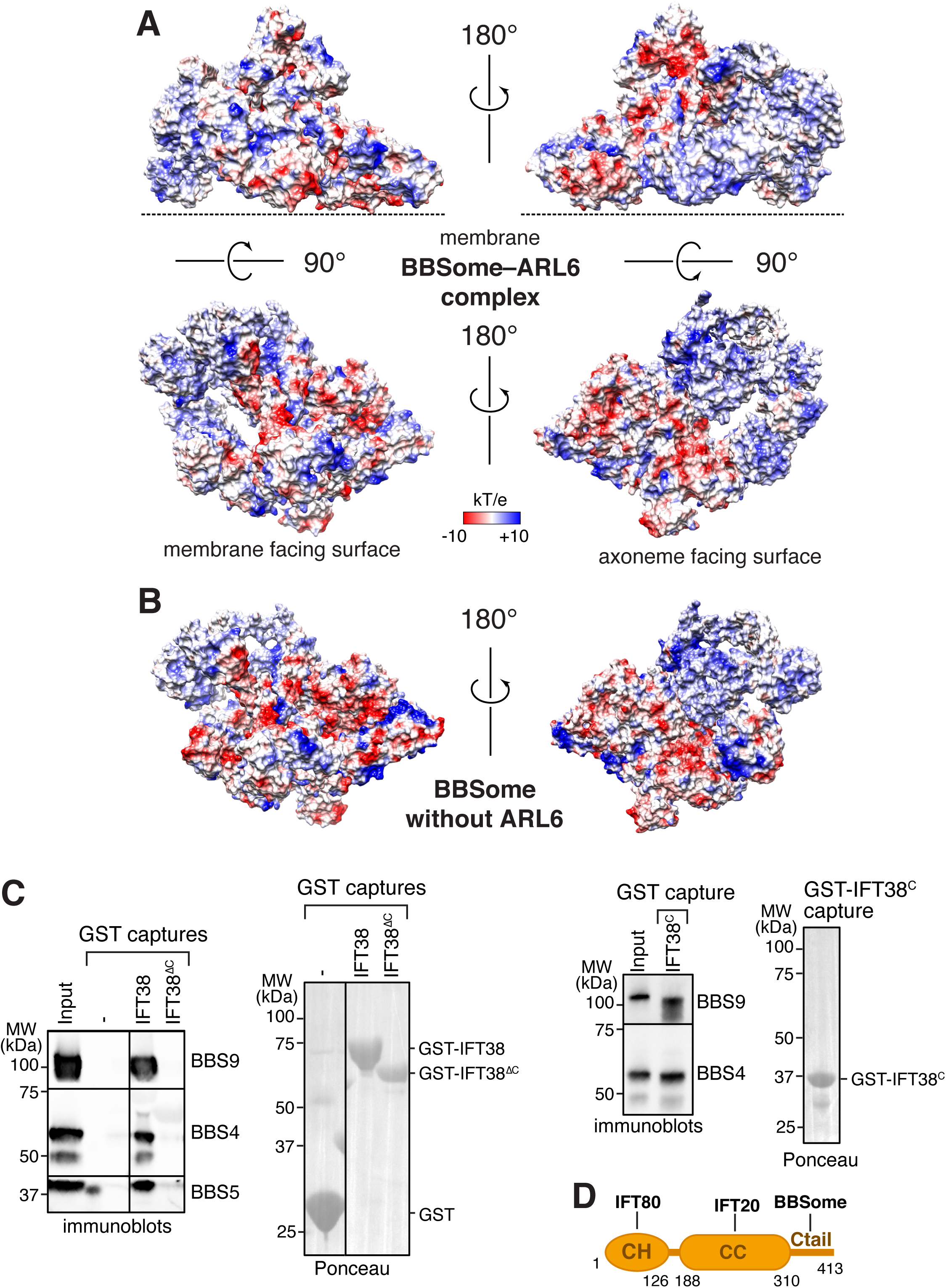
Proposed binding surfaces for IFT38 and membranes on the BBSome. **A**. Views of the electrostatic surface of the BBSome–ARL6^GTP^ complex. **B.** Views of the electrostatic surface of the BBSome alone. **C.** IFT38^Ctail^ is necessary and sufficient for BBSome binding. GST-capture assays were conducted with BBSome purified from bovine retina and GST fusions immobilized on glutathione sepharose. Bound material was eluted in SDS sample buffer. 2.5 input equivalents were loaded in the capture lanes. The BBSome was detected by immunoblotting and the GST fusions by Ponceau S staining. **D.** Diagram of the domain organization of IFT38. The calponin homology (CH) domain interacts with IFT80, the coiled-coil (CC) domain with IFT20, and the C-terminal tail (Ctail) with the BBSome.

**Supplement Figure 7.**
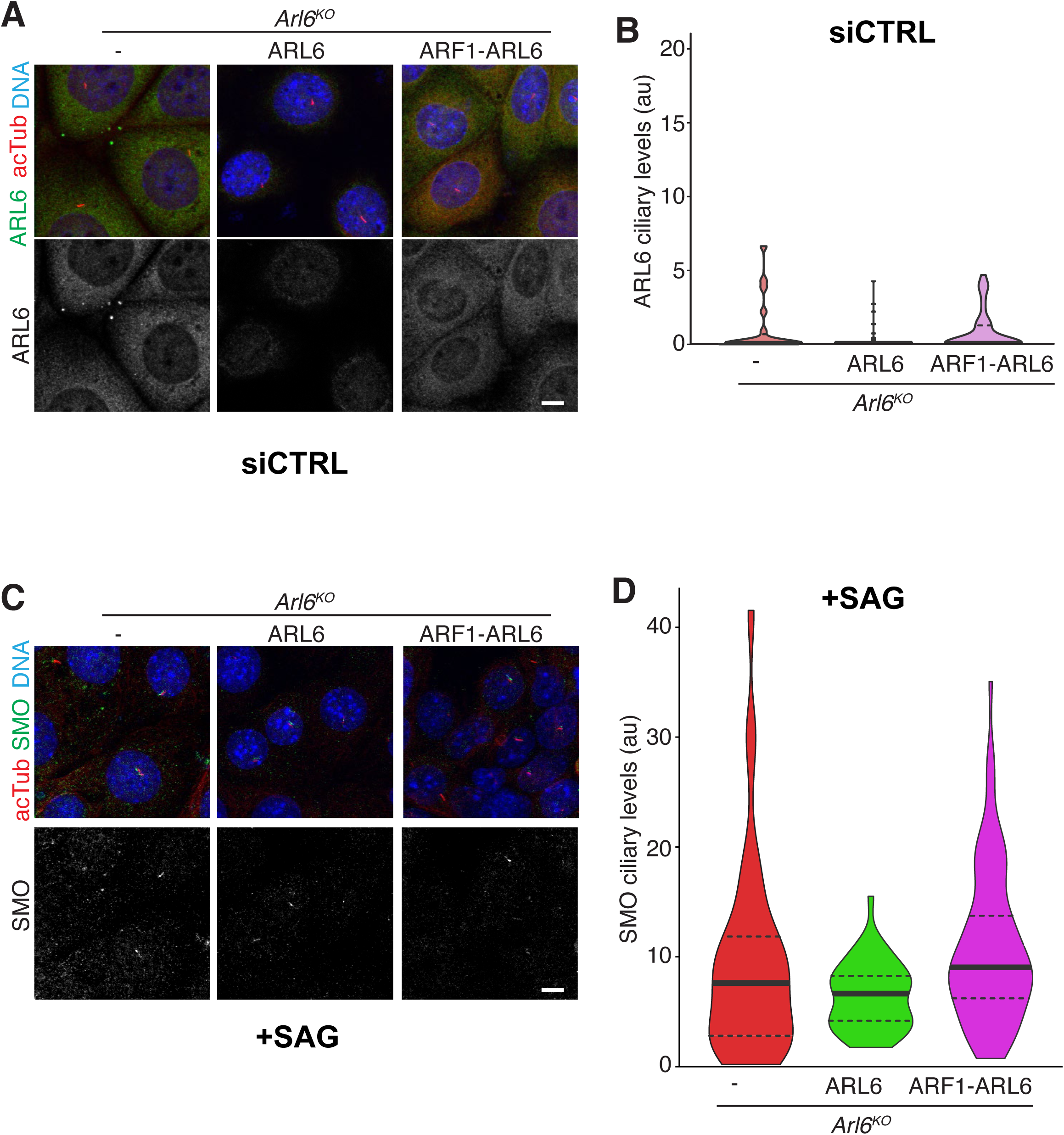
Additional characterization of the ARF1-ARL6 cell line. **A-B.** Control for Fig. 7D-E. In IMCD3 cells treated with control siRNA (siCTRL), the ciliary level of ARL6 remains undetectable by immunostaining. Cells were stained for ARL6, acetylated tubulin and DNA. Scale bar: 10 μm. **B.** The fluorescence intensity of the ARL6 channel in the cilium was measured in each cell line and the data are represented in a violin plot. **C-D.** Control for Fig. 7F-G. **C.** Cells were treated with Smoothened agonist (SAG) for 2h before fixation and staining for SMO, acetylated tubulin and DNA. Scale bar: 10 μm. **D.** The fluorescence intensity of the SMO channel in the cilium was measured in each cell line and the data are represented in a violin plot.

## SUPPLEMENTARY TABLES

**Table S1:** Cryo-EM data collection, refinement and modelling statistics.

**Table S2:** Fragments used in the BBS YTH array displayed in **Fig. 3D** and **S4C**.

**Table S3:** Sequences of helix 8 from known ciliary GPCRs.

## VIDEOS

**Video 1:** Morph of the BBSome structure from the unbound to the ARL6^GTP^ -bound conformation and back to unbound conformation. BBS1*^β^*^prop^ is blue, BBS4 is yellow and ARL6^GTP^ is magenta.

## MATERIALS AND METHODS

### Plasmid DNA

The plasmids for co-expressing ARL6 or ARF1-ARL6 together with ^AP^SSTR3^NG^ were built by introducing a second expression cassette into the pEF5ΔB/FRT/^AP^SSTR3^NG^ plasmid described in (Nager et al., 2017) and (Ye et al., 2018). The pEF5ΔB/FRT/^AP^SSTR3^NG^ plasmid expresses mouse SSTR3 fused to a biotinylation acceptor peptide (AP) at its extracellular N terminus and a mNeonGreen (NG) fluorescent protein at its intracellular C terminus. Expression of ^AP^SSTR3^NG^ is driven by the attenuated EF1α promoter (pEF1αΔ), which consists of a TATA-less EF1α promoter. This low-expression promoter enables reconstitution of the physiological exit kinetics of SSTR3 (Ye et al., 2018). Lastly, the pEF5ΔB/FRT backbone contains an FRT recombination site followed by a promoterless gene for blasticidin resistance to allow selection of recombinant cells in the recipient FlpIn cell lines. The second cassette was inserted into an NsiI site that had been introduced in the pEF5B/FRT plasmid between the ampicillin cassette and an EF1α promoter. The NsiI flanked cassettes for ARF1-ARL6 or ARL6 expression consist of an EF1α promoter followed by the sequence encoding ARF1-ARL6 or ARL6 and an HSV terminator.

The plasmids for SP6-driven *in vitro* transcription of individual BBSome subunits are based on pCS2+Myc6-DEST vectors and were described in (Jin et al., 2010). C-terminal truncations of BBS7 were generated by introducing stop codons using site-directed mutagenesis in pCS2+Myc6-BBS7. N-terminal truncations of BBS7 were assembled by PCR and Gateway recombination.

The plasmids for bacterial expression of SMO^ctail^, SMO^H8^ and IFT38 truncations are derivatives of pGEX6P1.

### Antibodies

Primary antibodies against the following proteins were used: actin (rabbit, Sigma-Aldrich, #A2066), cMyc (mouse, 9E10, Santa Cruz sc-40), acetylated tubulin (mouse, 6-11B-1, Sigma-Aldrich), ARL6 (rabbit, (Jin et al., 2010)), SMO (rabbit, gift from Kathryn Anderson, Memorial Sloan Kettering Cancer Center, New York, NY, (Ocbina et al., 2011)).

Secondary antibodies for immunofluorescence were: Cy3-conjugated donkey anti-rabbit IgG (7111-1655-152, Jackson Immunoresearch), Cy3-conjugated goat anti-mouse IgG2b (115-005-207, Jackson Immunoresearch), Cy5-conjugated goat anti-mouse IgG2b (115-175-207, Jackson Immunoresearch).

Secondary antibodies for immunoblotting were: HRP-conjugated goat anti-mouse IgG (115-035-003, Jackson Immunoresearch) and HRP-conjugated goat anti-rabbit IgG (111-035-003, Jackson Immunoresearch).

### Pharmacology

Pathway agonists were used in the following concentration: SAG (200 nM; ALX-270-426-M001; Enzo Life Sciences), somatostatin-14 (sst) (10 μM; ASR-003; Alomone Labs).

### Cell culture and generation of stable cell lines

All experiments were conducted in derivatives of the mouse IMCD3-FlpIn cell line. IMCD3-FlpIn cells were cultured in DMEM/F12, HEPES medium (11330-057, GIBCO) supplemented with 5% fetal bovine serum (FBS; Gemini Bio-Products), 100 U/mL penicillin-streptomycin (15140-122, GIBCO), and 2 mM L-glutamine (400-106, Gemini Bio-Products). Cell lines expressing ARL6, ARF1-ARL6, and/or ^AP^SSTR3^NG^ were generated using the Flp-In System (ThermoFisher Scientific) to ensure expression from a single genomic site and obtention of isogenic cell lines. *Arl6^-/-^* IMCD3-FlpIn cells (Liew et al., 2014) were transfected with pEF5ΔB/FRT derivatives along with a plasmid encoding the Flp-In recombinase (pOG44). Stable transformants were selected using 4 μg/mL Blasticidin (Invivogen) and individual colonies were picked. Clones with appropriate cilia morphology were selected for further study.

### Microscopy

For imaging of fixed specimens, 70,000 cells were seeded on acid-washed cover glasses (12 mm #1.5; 12-545-81; Thermo Fisher Scientific) in a 24-well plate. Cells were grown for 24 h and then starved for 20 h by shifting the cells to DMEM medium containing 0.2% FBS before pharmacological treatments. After treatment, cells were fixed with 4% paraformaldehyde in phosphate-buffered saline (PBS; 137 mM NaCl, 2.7 mM KCl, 10 mM Na_2_HPO_4_,1.8 mM KH_2_PO_4_) for 15 min at 37°C, then extracted in −20°C methanol for 5 min. The cells were permeabilized in PBS containing 0.1% Triton X-100, 5% normal donkey serum (017-000-121; Jackson ImmunoResearch Laboratories), and 3% bovine serum albumin (BP1605-100; Thermo Fisher Scientific) for 30 min. Subsequently, the permeabilized cells were immunostained by successive incubations with primary antibodies for 1 h, secondary antibodies for 30 min, and staining with Hoechst DNA dye, then mounted on slides using fluoromount-G (17984-25; Electron Microscopy Sciences).

Imaging of indirect immunofluorescence (i.e., all panels except Fig. 7D) was performed on a confocal microscope (LSM 800, Carl Zeiss Imaging) using a 63X/1.4NA objective and the 488, 555, 405 and 647 nm laser lines. Z-stacks of images with 0.2-µm separation were collected. Imaging of the direct fluorescence of cilia was performed using a DeltaVision system (Applied Precision) equipped with a PlanApo 60x/1.40 objective (Olympus), a pco.edge sCMOS camera and a solid-state illumination module (InsightSSI). For most experiments, cilia closest to the coverslip were imaged (ventral cilia) as these cilia often lay perpendicular to the objective. Z-stacks of images with 0.3-µm separation were collected.

The Z-stacks were imported to Fiji and maximal intensity projection were assembled. To measure the intensities of ciliary ARL6, SSTR3 and SMO, integrated fluorescence density was measured using Fiji and fluorescence of an adjacent area was subtracted as background.

### Sequence analysis

Helix 8 sequences were collected from GPCRdb (https://gpcrdb.org, (Pándy-Szekeres et al., 2018) and manually searched for BBSome-binding motifs. Ciliary GPCRs were collected from the literature (Badgandi et al., 2017; Berbari et al., 2008; Hilgendorf et al., 2019; Koemeter-Cox et al., 2014; Loktev and Jackson, 2013; Marley et al., 2013; Marley and von Zastrow, 2010; Mukhopadhyay et al., 2013; Omori et al., 2015; Siljee et al., 2018)

### Recombinant protein expression

N-terminally GST-tagged ARL6ΔN16[Q73L] was expressed in bacteria as described (Chou et al., 2019). GST-tagged SMO^Ctail^ and IFT38 protein fusions were expressed in Rosetta2(DE3)-pLysS cells grown in 2xYT medium (Millipore Sigma, Y2627) at 37°C until the optical density (OD) at 600 nm reached 0.6. Protein expression was then induced with 1 mM isopropyl *b*-D-1-thiogalactopyranoside (IPTG) at 18°C for 4 h (SMO^Ctail^) or with 0.2 mM IPTG at 18°C for 16 h (IFT38). Cells were resuspended in 4XT (200 mM Tris, pH 8.0, 800 mM NaCl, 1 mM DTT) with protease inhibitors (1 mM AEBSF, 0.8 μM Aprotinin, 15 μM E-64, 10 μg/mL Bestatin, 10 μg/mL Pepstatin A and 10 μg/mL Leupeptin) and lysed by sonication. The clarified lysates were loaded onto Glutathione Sepharose 4B resin (GE Healthcare) and proteins eluted with 50 mM reduced glutathione in buffer XT (50 mM Tris, pH 8.0, 200 mM NaCl, 1 mM DTT). Proteins were subsequently dialyzed against XT buffer with one change of buffer and flash frozen in liquid nitrogen after addition of 5% (w/v) glycerol.

### Purification of native BBSome

The BBSome was purified from bovine retina by ARL6^GTP^-affinity chromatography as described (Chou et al., 2019) and the sample was processed for cryo-EM the next day.

### GST-capture assays

GST pull-down assays were conducted by saturating 10 μL of Glutathione Sepharose 4B beads (GE #17075605) with GST fusions. Binding to purified BBSome was assessed by mixing beads with a 10 nM solution of pure BBSome made in 100 μL IB buffer (20 mM HEPES, pH 7.0, 5 mM MgCl_2_, 1 mM EDTA, 2% glycerol, 300 mM KOAc, 1 mM DTT, 0.2% Triton X-100) and incubating for 1 h at 4°C. After 4 washes with 200 μL IB buffer, elution was performed by boiling the beads in SDS sample buffer.

BBSome subunits and fragments thereof were translated *in vitro* from pCS2-Myc plasmids using the TNT SP6 Quick Coupled Transcription/Translation system (Promega L2080). 16 µL TNT SP6 Quick Master Mix, 2 µL Methionine (0.2 mM) and 2 µL DNA (0.2 µg/µL) were mixed and incubated at 30°C for 90 min. 20-µL reactions were diluted into 180 µL NSC250 buffer (25 mM Tris, pH 8.0, 250 mM KCl, 5 mM MgCl_2_, 0.5% CHAPS, 1 mM DTT), mixed with 10 μL glutathione beads saturated with GST fusions and rotated for 1 h at 4°C. After 4 washes with 200 μL NSC250 buffer, elution was performed by addition of 7.5 μg PreScission protease in 30 μL NSC250 buffer and incubation at 4°C overnight. The eluates were resolved by SDS-PAGE and analyzed by immunoblotting with anti-Myc antibody.

### Yeast two-hybrid assays

The coding DNA sequences (CDSs) for various fragments of BBSome subunits were either obtained in Gateway Entry vectors or amplified via PCR and transferred to pDONR221 by BP clonase recombination. The CDSs were shuttled to Y2H Gateway destination vectors bait pBTMcc24 (C-terminal bait), pBTM116D-9 (N-terminal bait), pCBDU (C-terminal prey), and pACT4 (N-terminal prey) by LR clonase recombination. Bait and prey vectors were introduced into either bait (L40ccU MATa) or prey (L40ccα MATα) yeast strains by lithium acetate transformation. Yeast were mated in a 96-well matrix format, using at least two independently transformed colonies to test each interaction. MATa and MATα yeast were mated on YPDA medium for 36-48 h at 30°C prior to diploid selection on medium lacking tryptophan and leucine. Diploids were incubated for 2 days at 30°C prior to transfer onto medium lacking tryptophan, leucine and histidine to select for positive growth of interacting constructs.

### Cryo-EM sample preparation and data collection

For BBSome alone, 3.5 μL of the peak fraction from a BBSome purification (0.4 - 0.6 mg/mL) was applied to glow-discharged R1.2/1.3 holey carbon copper grids (Quantifoil) covered with a thin homemade carbon film. The grids were blotted for 1 s at 4°C and 100% humidity, and plunged into liquid ethane using a Mark IV Vitrobot (Thermo Fisher Scientific). A cryo-EM dataset was collected on a 300-kV Titan Krios electron microscope (Thermo Fisher Scientific) equipped with a K2 Summit detector (Gatan) at a nominal magnification of 22,500x in super-resolution counting mode. After binning over 2×2 pixels, the calibrated pixel size was 1.3 Å on the specimen level.

Exposures of 10 s were dose-fractionated into 40 frames with a dose rate of 8 e^-^/pixel/s, resulting in a total dose of 80 e^-^/Å^2^. Data were collected with SerialEM (Mastronarde, 2005) and the used defocus range was from −1.5 μm to −3.0 μm.

For the BBSome–ARL6 complex, full-length ARL6 was incubated with GTP at a molar ratio of 1:20 for 1 h on ice, added to purified BBSome at a molar ratio of 5:1 and incubated for another hour on ice. 3.5 μL of the sample was applied to glow-discharged R1.2/1.3 holey carbon grids (Quantifoil Au or C-flat Cu). The grids were blotted for 3.5 s at 4°C and 100% humidity, and plunged into liquid ethane using a Mark IV Vitrobot. One cryo-EM dataset was collected on a 300-kV Titan Krios electron microscope equipped with a K2 Summit detector at a nominal magnification of 28,000x in super-resolution counting mode. After binning over 2×2 pixels, the calibrated pixel size was 1.0 Å on the specimen level. Exposures of 10 s were dose-fractionated into 40 frames with a dose rate of 7.52 e^-^/pixel/s, resulting in a total dose of 75.2 e^-^/Å^2^. A second dataset was collected on a 300-kV Titan Krios equipped with a K3 detector at a nominal magnification of 64,000x in super-resolution counting mode. After binning over 2×2 pixels, the calibrated pixel size was 1.08 Å on the specimen level. Exposures of 2 s were dose-fractionated into 50 frames with a dose rate of 29.99 e^-^/pixel/s, resulting in a total dose of 51.44 e^-^/Å^2^. Both datasets were collected with SerialEM and the defocus ranged from −1.5 μm to −2.5 μm.

### Cryo-EM data processing

The movie frames collected with the K2 detector were corrected with a gain reference. All movies were dose-weighted and motion-corrected with MotionCor2 (Zheng et al., 2017). The contrast transfer function (CTF) parameters were estimated with CTFFIND4 (Rohou and Grigorieff, 2015, p. 4). For micrographs collected with the K2 detector, particles were picked with Gautomatch (https://www.mrc-lmb.cam.ac.uk/kzhang/Gautomatch/); for those collected with the K3 detector, particles were picked with RELION 3.0 (Zivanov et al., 2018). Three projections from our previous cryo-EM map of the BBSome (EMD-7839) were used as templates for picking.

For BBSome alone, 2,218,320 particles were picked from 4,733 micrographs and subjected to 2D classification in RELION. Particles in classes that generated averages showing clear structural features were selected (770,345 particles) for *ab initio* 3D reconstruction of two models in cryoSPARC (Punjani et al., 2017). The map with clearer structural features and higher resolution was selected for heterogenous refinement in cryoSPARC, after which 560,777 particles were selected for further homogenous refinement. The output map was further refined in RELION by 3D refinement, CTF refinement and Bayesian polishing, resulting in a map at 3.6 Å resolution. The base and corkscrew modules of the BBSome, including BBS1, BBS4, BBS5, BBS8, BBS9 and BBS18, were well resolved, but density for the top lobe, containing BBS2 and BBS7, was weak. A focused refinement, masking out the BBS1 βprop and ins, BBS2 βprop, GAE and cc, and the BBS7 βprop, GAE, cc, pf and hp domains, yielded a map for the remainder of the BBSome at 3.44 Å resolution.

For the BBSome–ARL6 complex, data collected with the K2 camera yielded 228,487 particles from 1,031 micrographs of a Quantifoil Au grid and 192,243 particles from 911 images of a C-flat Cu grid. The particles from the two datasets were separately subjected to 2D classification in RELION, and particles from classes that generated averages showing clear structural features were combined, including 134,169 and 72,182 particles, respectively. Data collected with the K3 camera yielded 1,033,939 particles from 2,680 micrographs of the Quantifoil Au grid and 688,499 particles from 1,960 micrographs of the C-flat grid. After 2D classification, 185,332 and 154,503 particles, respectively, were selected. All selected particles were combined (546,186 particles in total) and subjected to 3D classification into 6 classes, using as reference the previously determined BBSome map (EMD-7839) filtered to 45-Å resolution. One of the resulting maps showed clear fine structural features (209,646 particles) and was subjected to 3D refinement, yielding a density map at 4.1-Å resolution. Refinement focused on the top lobe of the BBSome, including BBS2, BBS7, BBS1 βprop, and ARL6 yielded a map at 4.2-Å resolution. Refinement focused on the lower lobe of the BBSome including the remaining subunits yielded a map at 3.8-Å resolution. To improve the density for the GAE and pf domains of BBS2 and BBS7, a mask was generated for these domains and used for focused 3D classification into 4 classes without alignment. The resulting map showing the best structural features was selected for further refinement, which resulted in a map at 4.0-Å resolution, with improved density for the GAE and pf domains of BBS2 and BBS7.

The resolution was determined by Fourier shell correlation (FSC) of two independently refined half-maps using the 0.143 cut-off criterion (Rosenthal and Henderson, 2003). Local resolution was estimated from the two half-maps using the ResMap algorithm implemented in RELION. UCSF Chimera (Pettersen et al., 2004) was used to visualize density maps.

### Model building and refinement

Our previously published backbone model of the BBSome (Chou et al., 2019) was first placed into the density map using Chimera. All manual model building was performed with Coot (Emsley and Cowtan, 2004). BBS2^GAE^ and BBS7^GAE^ models were generated using SWISS-MODEL (Waterhouse et al., 2018), using the structure of BBS9^GAE^ as template. The generated models were then docked into the density map using Chimera, and trimmed in Coot. Due to the weak density of these areas in both maps, we only built secondary-structure fragments but not the connecting loops. A model for bovine ARL6 starting was generated with SWISS-MODEL, using the crystal structure of *Chlamydomonas reinhardtii* ARL6 (PDB ID: 40VN) as template. The model was then docked into the density map of the BBSome–ARL6 complex. The atomic models were refined using phenix.real_space_refine (Adams et al., 2010). Cryo-EM data collection, refinement and modelling statistics are summarized in Table S1.

## REFERENCES

Adams PD, Afonine PV, Bunkóczi G, Chen VB, Davis IW, Echols N, Headd JJ, Hung L-W, Kapral GJ, Grosse-Kunstleve RW, McCoy AJ, Moriarty NW, Oeffner R, Read RJ, Richardson DC, Richardson JS, Terwilliger TC, Zwart PH. 2010. PHENIX: a comprehensive Python-based system for macromolecular structure solution. Acta Crystallogr D Biol Crystallogr 66:213–221. doi:10.1107/S0907444909052925

Anvarian Z, Mykytyn K, Mukhopadhyay S, Pedersen LB, Christensen ST. 2019. Cellular signalling by primary cilia in development, organ function and disease. Nat Rev Nephrol. doi:10.1038/s41581-019-0116-9

Badgandi HB, Hwang S, Shimada IS, Loriot E, Mukhopadhyay S. 2017. Tubby family proteins are adapters for ciliary trafficking of integral membrane proteins. J Cell Biol 216:743–760. doi:10.1083/jcb.201607095

Bangs F, Anderson KV. 2017. Primary cilia and mammalian Hedgehog signaling. Cold Spring Harb Perspect Biol 9:a028175. doi:10.1101/cshperspect.a028175

Berbari NF, Johnson AD, Lewis JS, Askwith CC, Mykytyn K. 2008. Identification of Ciliary Localization Sequences within the Third Intracellular Loop of G Protein-coupled Receptors. Mol Biol Cell 19:1540–1547. doi:10.1091/mbc.E07-09-0942

Bhatia VK, Hatzakis NS, Stamou D. 2010. A unifying mechanism accounts for sensing of membrane curvature by BAR domains, amphipathic helices and membrane-anchored proteins. Semin Cell Dev Biol 21:381–390. doi:10.1016/j.semcdb.2009.12.004

Bi X, Mancias JD, Goldberg J. 2007. Insights into COPII coat nucleation from the structure of Sec23*Sar1 complexed with the active fragment of Sec31. Dev Cell 13:635–645. doi:10.1016/j.devcel.2007.10.006

Boldt K, van Reeuwijk J, Lu Q, Koutroumpas K, Nguyen T-MT, Texier Y, van Beersum SEC, Horn N, Willer JR, Mans DA, Dougherty G, Lamers IJC, Coene KLM, Arts HH, Betts MJ, Beyer T, Bolat E, Gloeckner CJ, Haidari K, Hetterschijt L, Iaconis D, Jenkins D, Klose F, Knapp B, Latour B, Letteboer SJF, Marcelis CL, Mitic D, Morleo M, Oud MM, Riemersma M, Rix S, Terhal PA, Toedt G, van Dam TJP, de Vrieze E, Wissinger Y, Wu KM, Apic G, Beales PL, Blacque OE, Gibson TJ, Huynen MA, Katsanis N, Kremer H, Omran H, van Wijk E, Wolfrum U, Kepes F, Davis EE, Franco B, Giles RH, Ueffing M, Russell RB, Roepman R, UK10K Rare Diseases Group. 2016. An organelle-specific protein landscape identifies novel diseases and molecular mechanisms. Nat Commun 7:11491. doi:10.1038/ncomms11491

Breslow D, Koslover EF, Seydel F, Spakowitz AJ, Nachury MV. 2013. An in vitro assay for entry into cilia reveals unique properties of the soluble diffusion barrier. J Cell Biol 203:129– 147. doi:10.1083/jcb.201212024

Byrne EFX, Sircar R, Miller PS, Hedger G, Luchetti G, Nachtergaele S, Tully MD, Mydock-McGrane L, Covey DF, Rambo RP, Sansom MSP, Newstead S, Rohatgi R, Siebold C. 2016. Structural basis of Smoothened regulation by its extracellular domains. Nature 535:517–522. doi:10.1038/nature18934

Chadha A, Volland S, Baliaouri NV, Tran EM, Williams DS. 2019. The route of the visual receptor rhodopsin along the cilium. J Cell Sci 132. doi:10.1242/jcs.229526

Cherfils J. 2014. Arf GTPases and their effectors: assembling multivalent membrane-binding platforms. Curr Opin Struct Biol 29:67–76. doi:10.1016/j.sbi.2014.09.007

Chou H-T, Apelt L, Farrell DP, White SR, Woodsmith J, Svetlov V, Goldstein JS, Nager AR, Li Z, Muller J, Dollfus H, Nudler E, Stelzl U, DiMaio F, Nachury MV, Walz T. 2019. The Molecular Architecture of Native BBSome Obtained by an Integrated Structural Approach. Struct Lond Engl 1993 27:1384–1394.e4. doi:10.1016/j.str.2019.06.006

Deshpande I, Liang J, Hedeen D, Roberts KJ, Zhang Y, Ha B, Latorraca NR, Faust B, Dror RO, Beachy PA, Myers BR, Manglik A. 2019. Smoothened stimulation by membrane sterols drives Hedgehog pathway activity. Nature 571:284–288. doi:10.1038/s41586-019-1355-4

Dodonova SO, Aderhold P, Kopp J, Ganeva I, Röhling S, Hagen WJH, Sinning I, Wieland F, Briggs JAG. 2017. 9 Å structure of the COPI coat reveals that the Arf1 GTPase occupies two contrasting molecular environments. eLife 6:352. doi:10.7554/eLife.26691

Drin G, Antonny B. 2010. Amphipathic helices and membrane curvature. FEBS Lett 584:1840– 1847. doi:10.1016/j.febslet.2009.10.022

Eguether T, San Agustin JT, Keady BT, Jonassen JA, Liang Y, Francis R, Tobita K, Johnson CA, Abdelhamed ZA, Lo CW, Pazour GJ. 2014. IFT27 links the BBSome to IFT for maintenance of the ciliary signaling compartment. Dev Cell 31:279–290. doi:10.1016/j.devcel.2014.09.011

Emsley P, Cowtan K. 2004. Coot: model-building tools for molecular graphics. Acta Crystallogr D Biol Crystallogr 60:2126–2132. doi:10.1107/S0907444904019158

Faini M, Beck R, Wieland FT, Briggs JAG. 2013. Vesicle coats: structure, function, and general principles of assembly. Trends Cell Biol 23:279–288. doi:10.1016/j.tcb.2013.01.005

Fisch C, Dupuis Williams P. 2011. Ultrastructure of cilia and flagella - back to the future! Biol Cell Auspices Eur Cell Biol Organ 103:249–270. doi:10.1042/BC20100139

Gallagher BC. 1980. Primary cilia of the corneal endothelium. Am J Anat 159:475–484. doi:10.1002/aja.1001590410

Garcia-Gonzalo FR, Reiter JF. 2017. Open Sesame: how transition fibers and the transition zone control ciliary composition. Cold Spring Harb Perspect Biol 9:a028134. doi:10.1101/cshperspect.a028134

Gillingham AK, Munro S. 2007. The small G proteins of the Arf family and their regulators. Annu Rev Cell Dev Biol 23:579–611. doi:10.1146/annurev.cellbio.23.090506.123209

Goetz SC, Bangs F, Barrington CL, Katsanis N, Anderson KV. 2017. The Meckel syndrome-associated protein MKS1 functionally interacts with components of the BBSome and IFT complexes to mediate ciliary trafficking and hedgehog signaling. PLoS ONE 12:e0173399. doi:10.1371/journal.pone.0173399

Green JA, Schmid CL, Bley E, Monsma PC, Brown A, Bohn LM, Mykytyn K. 2015. Recruitment of β-Arrestin into Neuronal Cilia Modulates Somatostatin Receptor Subtype 3 Ciliary Localization. Mol Cell Biol 36:223–235. doi:10.1128/MCB.00765-15

Hatzakis NS, Bhatia VK, Larsen J, Madsen KL, Bolinger P-Y, Kunding AH, Castillo J, Gether U, Hedegård P, Stamou D. 2009. How curved membranes recruit amphipathic helices and protein anchoring motifs. Nat Chem Biol 5:835–841. doi:10.1038/nchembio.213

Hilgendorf KI, Johnson CT, Mezger A, Rice SL, Norris AM, Demeter J, Greenleaf WJ, Reiter JF, Kopinke D, Jackson PK. 2019. Omega-3 Fatty Acids Activate Ciliary FFAR4 to Control Adipogenesis. Cell 179:1289–1305.e21. doi:10.1016/j.cell.2019.11.005

Huang P, Zheng S, Wierbowski BM, Kim Y, Nedelcu D, Aravena L, Liu J, Kruse AC, Salic A. 2018. Structural Basis of Smoothened Activation in Hedgehog Signaling. Cell 174:312–324.e16. doi:10.1016/j.cell.2018.04.029

Isakoff SJ, Cardozo T, Andreev J, Li Z, Ferguson KM, Abagyan R, Lemmon MA, Aronheim A, Skolnik EY. 1998. Identification and analysis of PH domain-containing targets of phosphatidylinositol 3-kinase using a novel in vivo assay in yeast. EMBO J 17:5374–5387. doi:10.1093/emboj/17.18.5374

Jackson LP, Kelly BT, McCoy AJ, Gaffry T, James LC, Collins BM, Höning S, Evans PR, Owen DJ. 2010. A large-scale conformational change couples membrane recruitment to cargo binding in the AP2 clathrin adaptor complex. Cell 141:1220–1229. doi:10.1016/j.cell.2010.05.006

Jana SC, Mendonça S, Machado P, Werner S, Rocha J, Pereira A, Maiato H, Bettencourt-Dias M. 2018. Differential regulation of transition zone and centriole proteins contributes to ciliary base diversity. Nat Cell Biol 20:928–941. doi:10.1038/s41556-018-0132-1

Jian X, Tang W-K, Zhai P, Roy NS, Luo R, Gruschus JM, Yohe ME, Chen P-W, Li Y, Byrd RA, Xia D, Randazzo PA. 2015. Molecular Basis for Cooperative Binding of Anionic Phospholipids to the PH Domain of the Arf GAP ASAP1. Structure 23:1977–1988. doi:10.1016/j.str.2015.08.008

Jin H, White SR, Shida T, Schulz S, Aguiar M, Gygi SP, Bazan JF, Nachury MV. 2010. The conserved Bardet-Biedl syndrome proteins assemble a coat that traffics membrane proteins to cilia. Cell 141:1208–1219. doi:10.1016/j.cell.2010.05.015

Kamiya R, Witman GB. 1984. Submicromolar levels of calcium control the balance of beating between the two flagella in demembranated models of Chlamydomonas. J Cell Biol 98:97–107.

Klink BU, Gatsogiannis C, Hofnagel O, Wittinghofer A, Raunser S. 2020. Structure of the human BBSome core complex. eLife 9. doi:10.7554/eLife.53910

Klink BU, Zent E, Juneja P, Kuhlee A, Raunser S, Wittinghofer A. 2017. A recombinant BBSome core complex and how it interacts with ciliary cargo. eLife 6. doi:10.7554/eLife.27434

Koemeter-Cox AI, Sherwood TW, Green JA, Steiner RA, Berbari NF, Yoder BK, Kauffman AS, Monsma PC, Brown A, Askwith CC, Mykytyn K. 2014. Primary cilia enhance kisspeptin receptor signaling on gonadotropin-releasing hormone neurons. Proc Natl Acad Sci U S A 111:10335–10340. doi:10.1073/pnas.1403286111

Krishna AG, Menon ST, Terry TJ, Sakmar TP. 2002. Evidence that helix 8 of rhodopsin acts as a membrane-dependent conformational switch. Biochemistry 41:8298–8309. doi:10.1021/bi025534m

Larsen JB, Jensen MB, Bhatia VK, Pedersen SL, Bjørnholm T, Iversen L, Uline M, Szleifer I, Jensen KJ, Hatzakis NS, Stamou D. 2015. Membrane curvature enables N-Ras lipid anchor sorting to liquid-ordered membrane phases. Nat Chem Biol 11:192–194. doi:10.1038/nchembio.1733

Li Z-L. 2018. Molecular dynamics simulations of membrane deformation induced by amphiphilic helices of Epsin, Sar1p, and Arf1. Chin Phys B 27:038703. doi:10.1088/1674-1056/27/3/038703

Liew GM, Ye F, Nager AR, Murphy JP, Lee JS, Aguiar M, Breslow D, Gygi SP, Nachury MV. 2014. The intraflagellar transport protein IFT27 promotes BBSome exit from cilia through the GTPase ARL6/BBS3. Dev Cell 31:265–278. doi:10.1016/j.devcel.2014.09.004

Liu P, Lechtreck KF. 2018. The Bardet–Biedl syndrome protein complex is an adapter expanding the cargo range of intraflagellar transport trains for ciliary export. Proc Natl Acad Sci 115:E934–E943. doi:10.1073/pnas.1713226115

Loktev AV, Jackson PK. 2013. Neuropeptide Y family receptors traffic via the Bardet-Biedl syndrome pathway to signal in neuronal primary cilia. Cell Rep 5:1316–1329. doi:10.1016/j.celrep.2013.11.011

Makyio H, Ohgi M, Takei T, Takahashi S, Takatsu H, Katoh Y, Hanai A, Ueda T, Kanaho Y, Xie Y, Shin H-W, Kamikubo H, Kataoka M, Kawasaki M, Kato R, Wakatsuki S, Nakayama K. 2012. Structural basis for Arf6–MKLP1 complex formation on the Flemming body responsible for cytokinesis. EMBO J 31:2590–2603. doi:10.1038/emboj.2012.89

Manneville J-B, Casella J-F, Ambroggio E, Gounon P, Bertherat J, Bassereau P, Cartaud J, Antonny B, Goud B. 2008. COPI coat assembly occurs on liquid-disordered domains and the associated membrane deformations are limited by membrane tension. Proc Natl Acad Sci U S A 105:16946–16951. doi:10.1073/pnas.0807102105

Marley A, Choy RW-Y, von Zastrow M. 2013. GPR88 reveals a discrete function of primary cilia as selective insulators of GPCR cross-talk. PLoS ONE 8:e70857. doi:10.1371/journal.pone.0070857

Marley A, von Zastrow M. 2010. DISC1 regulates primary cilia that display specific dopamine receptors. PLoS ONE 5:e10902. doi:10.1371/journal.pone.0010902

Mastronarde DN. 2005. Automated electron microscope tomography using robust prediction of specimen movements. J Struct Biol 152:36–51. doi:10.1016/j.jsb.2005.07.007

Milenkovic L, Milenkovic L, Scott MP, Scott MP, Rohatgi R. 2009. Lateral transport of Smoothened from the plasma membrane to the membrane of the cilium. J Cell Biol 187:365–374. doi:10.1083/jcb.200907126

Milenkovic L, Weiss LE, Yoon J, Roth TL, Su YS, Sahl SJ, Scott MP, Moerner WE. 2015. Single-molecule imaging of Hedgehog pathway protein Smoothened in primary cilia reveals binding events regulated by Patched1. Proc Natl Acad Sci U A 112:8320–8325. doi:10.1073/pnas.1510094112

Miller SE, Mathiasen S, Bright NA, Pierre F, Kelly BT, Kladt N, Schauss A, Merrifield CJ, Stamou D, Höning S, Owen DJ. 2015. CALM regulates clathrin-coated vesicle size and maturation by directly sensing and driving membrane curvature. Dev Cell 33:163–175. doi:10.1016/j.devcel.2015.03.002

Mourão A, Nager AR, Nachury MV, Lorentzen E. 2014. Structural basis for membrane targeting of the BBSome by ARL6. Nat Struct Mol Biol 21:1035–1041. doi:10.1038/nsmb.2920

Mukhopadhyay S, Wen X, Ratti N, Loktev A, Rangell L, Scales SJ, Jackson PK. 2013. The ciliary G-protein-coupled receptor Gpr161 negatively regulates the Sonic hedgehog pathway via cAMP signaling. Cell 152:210–223. doi:10.1016/j.cell.2012.12.026

Nachury MV. 2018. The molecular machines that traffic signaling receptors into and out of cilia. Curr Opin Cell Biol 51:124–131. doi:10.1016/j.ceb.2018.03.004

Nachury MV, Loktev AV, Zhang Q, Westlake CJ, Peränen J, Merdes A, Slusarski DC, Scheller RH, Bazan JF, Sheffield VC, Jackson PK. 2007. A core complex of BBS proteins cooperates with the GTPase Rab8 to promote ciliary membrane biogenesis. Cell 129:1201–1213. doi:10.1016/j.cell.2007.03.053

Nachury MV, Mick DU. 2019. Establishing and regulating the composition of cilia for signal transduction. Nat Rev Mol Cell Biol 20:389–405. doi:10.1038/s41580-019-0116-4

Nager AR, Goldstein JS, Herranz-Pérez V, Portran D, Ye F, García-Verdugo JM, Nachury MV. 2017. An Actin Network Dispatches Ciliary GPCRs into Extracellular Vesicles to Modulate Signaling. Cell 168:252–263.e14. doi:10.1016/j.cell.2016.11.036

Nozaki S, Castro Araya RF, Katoh Y, Nakayama K. 2019. Requirement of IFT-B-BBSome complex interaction in export of GPR161 from cilia. Biol Open 8. doi:10.1242/bio.043786

Ocbina PJR, Eggenschwiler JT, Moskowitz I, Anderson KV. 2011. Complex interactions between genes controlling trafficking in primary cilia. Nat Genet 43:547–553. doi:10.1038/ng.832

Omori Y, Chaya T, Yoshida S, Irie S, Tsujii T, Furukawa T. 2015. Identification of G Protein-Coupled Receptors (GPCRs) in Primary Cilia and Their Possible Involvement in Body Weight Control. PLoS ONE 10:e0128422. doi:10.1371/journal.pone.0128422

Paczkowski JE, Fromme JC. 2014. Structural basis for membrane binding and remodeling by the exomer secretory vesicle cargo adaptor. Dev Cell 30:610–624. doi:10.1016/j.devcel.2014.07.014

Pándy-Szekeres G, Munk C, Tsonkov TM, Mordalski S, Harpsøe K, Hauser AS, Bojarski AJ, Gloriam DE. 2018. GPCRdb in 2018: adding GPCR structure models and ligands. Nucleic Acids Res 46:D440–D446. doi:10.1093/nar/gkx1109

Panic B, Perisic O, Veprintsev DB, Williams RL, Munro S. 2003. Structural Basis for Arl1-Dependent Targeting of Homodimeric GRIP Domains to the Golgi Apparatus. Mol Cell 12:863–874. doi:10.1016/S1097-2765(03)00356-3

Pettersen EF, Goddard TD, Huang CC, Couch GS, Greenblatt DM, Meng EC, Ferrin TE. 2004. UCSF Chimera–a visualization system for exploratory research and analysis. J Comput Chem 25:1605–1612. doi:10.1002/jcc.20084

Piscitelli CL, Kean J, Graaf C de, Deupi X. 2015. A molecular pharmacologist’s guide to G protein–coupled receptor crystallography. Mol Pharmacol 88:536–551. doi:10.1124/mol.115.099663

Punjani A, Rubinstein JL, Fleet DJ, Brubaker MA. 2017. cryoSPARC: algorithms for rapid unsupervised cryo-EM structure determination. Nat Methods 14:290–296. doi:10.1038/nmeth.4169

Qi X, Liu H, Thompson B, McDonald J, Zhang C, Li X. 2019. Cryo-EM structure of oxysterol-bound human Smoothened coupled to a heterotrimeric Gi. Nature 571:279–283. doi:10.1038/s41586-019-1286-0

Ren X, Farías GG, Canagarajah BJ, Bonifacino JS, Hurley JH. 2013. Structural basis for recruitment and activation of the AP-1 clathrin adaptor complex by Arf1. Cell 152:755– 767. doi:10.1016/j.cell.2012.12.042

Ringo DL. 1967. Flagellar motion and fine structure of the flagellar apparatus in Chlamydomonas. J Cell Biol 33:543–571.

Rohou A, Grigorieff N. 2015. CTFFIND4: Fast and accurate defocus estimation from electron micrographs. J Struct Biol 192:216–221. doi:10.1016/j.jsb.2015.08.008

Rosenthal PB, Henderson R. 2003. Optimal determination of particle orientation, absolute hand, and contrast loss in single-particle electron cryomicroscopy. J Mol Biol 333:721– 745. doi:10.1016/j.jmb.2003.07.013

Sato T, Kawasaki T, Mine S, Matsumura H. 2016. Functional role of the C-terminal smphipathic helix 8 of olfactory receptors and other G protein-coupled receptors. Int J Mol Sci 17:1930. doi:10.3390/ijms17111930

Schmidt HB, Görlich D. 2016. Transport Selectivity of Nuclear Pores, Phase Separation, and Membraneless Organelles. Trends Biochem Sci, Special Issue: 40 Years of TiBS 41:46–61. doi:10.1016/j.tibs.2015.11.001

Schou KB, Mogensen JB, Morthorst SK, Nielsen BS, Aleliunaite A, Serra-Marques A, Fürstenberg N, Saunier S, Bizet AA, Veland IR, Akhmanova A, Christensen ST, Pedersen LB. 2017. KIF13B establishes a CAV1-enriched microdomain at the ciliary transition zone to promote Sonic hedgehog signalling. Nat Commun 8:14177. doi:10.1038/ncomms14177

Seelig J. 2004. Thermodynamics of lipid-peptide interactions. Biochim Biophys Acta 1666:40–50. doi:10.1016/j.bbamem.2004.08.004

Seo S, Zhang Q, Bugge K, Breslow D, Searby CC, Nachury MV, Sheffield VC. 2011. A novel protein LZTFL1 regulates ciliary trafficking of the BBSome and Smoothened. PLoS Genet 7:e1002358. doi:10.1371/journal.pgen.1002358

Siljee JE, Wang Y, Bernard AA, Ersoy BA, Zhang S, Marley A, Von Zastrow M, Reiter JF, Vaisse C. 2018. Subcellular localization of MC4R with ADCY3 at neuronal primary cilia underlies a common pathway for genetic predisposition to obesity. Nat Genet 50:180–185. doi:10.1038/s41588-017-0020-9

Singh SK, Gui M, Koh F, Yip MC, Brown A. 2020. Structure and activation mechanism of the BBSome membrane protein trafficking complex. eLife 9. doi:10.7554/eLife.53322

Sztul E, Chen P-W, Casanova JE, Cherfils J, Dacks JB, Lambright DG, Lee F-JS, Randazzo PA, Santy LC, Schürmann A, Wilhelmi I, Yohe ME, Kahn RA. 2019. ARF GTPases and their GEFs and GAPs: concepts and challenges. Mol Biol Cell 30:1249–1271. doi:10.1091/mbc.E18-12-0820

Trépout S, Tassin A-M, Marco S, Bastin P. 2018. STEM tomography analysis of the trypanosome transition zone. J Struct Biol 202:51–60. doi:10.1016/j.jsb.2017.12.005

Vieira OV, Gaus K, Verkade P, Fullekrug J, Vaz WLC, Simons K. 2006. FAPP2, cilium formation, and compartmentalization of the apical membrane in polarized Madin-Darby canine kidney (MDCK) cells. Proc Natl Acad Sci U S A 103:18556–18561. doi:10.1073/pnas.0608291103

Vonkova I, Saliba A-E, Deghou S, Anand K, Ceschia S, Doerks T, Galih A, Kugler KG, Maeda K, Rybin V, van Noort V, Ellenberg J, Bork P, Gavin A-C. 2015. Lipid Cooperativity as a General Membrane-Recruitment Principle for PH Domains. Cell Rep 12:1519–1530. doi:10.1016/j.celrep.2015.07.054

Wang C, Wu H, Evron T, Vardy E, Han GW, Huang X-P, Hufeisen SJ, Mangano TJ, Urban DJ, Katritch V, Cherezov V, Caron MG, Roth BL, Stevens RC. 2014. Structural basis for Smoothened receptor modulation and chemoresistance to anticancer drugs. Nat Commun 5:4355. doi:10.1038/ncomms5355

Wang C, Wu H, Katritch V, Han GW, Huang X-P, Liu W, Siu FY, Roth BL, Cherezov V, Stevens RC. 2013. Structure of the human smoothened receptor bound to an antitumour agent. Nature 497:338–343. doi:10.1038/nature12167

Waterhouse A, Bertoni M, Bienert S, Studer G, Tauriello G, Gumienny R, Heer FT, de Beer TAP, Rempfer C, Bordoli L, Lepore R, Schwede T. 2018. SWISS-MODEL: homology modelling of protein structures and complexes. Nucleic Acids Res 46:W296–W303. doi:10.1093/nar/gky427

Weierstall U, James D, Wang C, White TA, Wang D, Liu W, Spence JCH, Bruce Doak R, Nelson G, Fromme P, Fromme R, Grotjohann I, Kupitz C, Zatsepin NA, Liu H, Basu S, Wacker D, Han GW, Katritch V, Boutet S, Messerschmidt M, Williams GJ, Koglin JE, Marvin Seibert M, Klinker M, Gati C, Shoeman RL, Barty A, Chapman HN, Kirian RA, Beyerlein KR, Stevens RC, Li D, Shah STA, Howe N, Caffrey M, Cherezov V. 2014. Lipidic cubic phase injector facilitates membrane protein serial femtosecond crystallography. Nat Commun 5:3309. doi:10.1038/ncomms4309

Wiens CJ, Tong Y, Esmail MA, Oh E, Gerdes JM, Wang J, Tempel W, Rattner JB, Katsanis N, Park H-W, Leroux MR. 2010. Bardet-Biedl syndrome-associated small GTPase ARL6 (BBS3) functions at or near the ciliary gate and modulates Wnt signaling. J Biol Chem 285:16218–16230. doi:10.1074/jbc.M109.070953

Wingfield JL, Lechtreck K-F, Lorentzen E. 2018. Trafficking of ciliary membrane proteins by the intraflagellar transport/BBSome machinery. Essays Biochem 62:753–763. doi:10.1042/EBC20180030

Woodsmith J, Apelt L, Casado-Medrano V, Özkan Z, Timmermann B, Stelzl U. 2017. Protein interaction perturbation profiling at amino-acid resolution. Nat Methods 14:1213–1221. doi:10.1038/nmeth.4464

Ye F, Nager AR, Nachury MV. 2018. BBSome trains remove activated GPCRs from cilia by enabling passage through the transition zone. J Cell Biol 217:1847–1868. doi:10.1083/jcb.201709041

Yu JW, Mendrola JM, Audhya A, Singh S, Keleti D, DeWald DB, Murray D, Emr SD, Lemmon MA. 2004. Genome-wide analysis of membrane targeting by S. cerevisiae pleckstrin homology domains. Mol Cell 13:677–688. doi:10.1016/s1097-2765(04)00083-8

Yu X, Breitman M, Goldberg J. 2012. A Structure-Based Mechanism for Arf1-Dependent Recruitment of Coatomer to Membranes. Cell 148:530–542. doi:10.1016/j.cell.2012.01.015

Zhang Q, Nishimura D, Seo S, Vogel T, Morgan DA, Searby C, Bugge K, Stone EM, Rahmouni K, Sheffield VC. 2011. Bardet-Biedl syndrome 3 (Bbs3) knockout mouse model reveals common BBS-associated phenotypes and Bbs3 unique phenotypes. Proc Natl Acad Sci U A 108:20678–20683. doi:10.1073/pnas.1113220108

Zhang X, Zhao F, Wu Y, Yang J, Han GW, Zhao S, Ishchenko A, Ye L, Lin X, Ding K, Dharmarajan V, Griffin PR, Gati C, Nelson G, Hunter MS, Hanson MA, Cherezov V, Stevens RC, Tan W, Tao H, Xu F. 2017. Crystal structure of a multi-domain human smoothened receptor in complex with a super stabilizing ligand. Nat Commun 8:15383. doi:10.1038/ncomms15383

Zheng SQ, Palovcak E, Armache J-P, Verba KA, Cheng Y, Agard DA. 2017. MotionCor2: anisotropic correction of beam-induced motion for improved cryo-electron microscopy. Nat Methods 14:331–332. doi:10.1038/nmeth.4193

Zivanov J, Nakane T, Forsberg BO, Kimanius D, Hagen WJ, Lindahl E, Scheres SH. 2018. New tools for automated high-resolution cryo-EM structure determination in RELION-3. eLife 7. doi:10.7554/eLife.42166

